# Multi-signal regulation of the GSK-3β homolog Rim11 governs meiosis entry in yeast

**DOI:** 10.1101/2023.09.21.558844

**Authors:** Johanna Kociemba, Andreas Christ Sølvsten Jørgensen, Nika Tadic, Anthony Harris, Theodora Sideri, Wei Yee Chan, Fairouz Ibrahim, Elcin Unal, Mark Skehel, Vahid Shahrezaei, Orlando Arguello-Miranda, Folkert J. van Werven

## Abstract

Starvation of budding yeast diploid cells induces the cell-fate program that drives meiosis and spore formation. Transcription activation of early meiotic genes (EMGs) requires the transcription activator Ime1, its DNA-binding partner Ume6, and GSK-3β kinase Rim11. Phosphorylation of Ume6 by Rim11 is key for EMG activation. We report that Rim11 integrates multiple input signals to control Ume6 phosphorylation and EMG transcription. Under nutrient-rich conditions PKA represses Rim11 to low levels while TORC1 keeps Rim11 localized to the cytoplasm. Inhibiting PKA and TORC1 induces Rim11 expression and nuclear localization. Remarkably, nuclear Rim11 is required, but not sufficient, for Rim11-dependent Ume6 phosphorylation. Additionally, Ime1 is an essential anchor protein for phosphorylating Ume6. Subsequently, Ume6-Ime1 coactivator complexes form that drive EMG transcription. Our results demonstrate how varied signalling inputs (PKA/TORC1/Ime1) integrated by Rim11 determine EMG expression and entry into meiosis. We propose that the signalling-regulatory network described here generates robustness in cell-fate control.

## Introduction

How cells decide to take a new fate is critical for development. The underlying signalling and regulatory programs ensure that fate decisions are taken at the correct time and space. Mis-regulated cell differentiation can be detrimental for development and cause disease pathologies. Understanding the molecular mechanisms including regulatory controls of underlying cell fate decisions remain a longstanding aim in molecular, cell and developmental biology.

In budding yeast, cells can undergo a critical cell fate decision called sporulation or gametogenesis during which a diploid cell produces four haploid spores. During sporulation diploid cells undergo premeiotic DNA replication, two consecutive chromosome divisions called meiosis, and subsequently packaging into haploid spores ^1^. The sporulation program is essential for reshuffling of genetic information and is critical for survival of yeasts during harsh environmental conditions.

The regulatory programme that controls entry into meiosis is set in motion by a master regulator called Ime1^2–4^. Cells lacking *IME1* will not enter meiosis nor form spores. *IME1* transcription is highly regulated through its complex promoter. The *IME1* promoter is repressed in cells harbouring a single mating type, which ensures that haploid cells cannot enter meiosis^5,6^. Moreover, TORC1 and PKA signalling keep the Tup1-Cyc8 co-repressor at the *IME1* promoter in a repressed state in nutrient rich conditions^7,8^. When diploid cells are starved, TORC1 and PKA signalling to *IME1* promoter is not active, Tup1-Cyc8 is relieved from the *IME1* promoter allowing for *IME1* transcription and consequently cells enter meiosis.

Ime1 is not sufficient to drive entry into meiosis. Two additional proteins, Ume6 and Rim11, are part of the regulatory circuit that activates transcription of the early meiotic genes (EMGs)^9–12^. Ume6 is the DNA binding component, while Ime1 harbours the transcriptional activation domain function. Under nutrient rich conditions Ume6 is a repressor of EMGs, by recruiting the co-repressor and histone deacetylase complex Sin3-Rpd3L^9,13–15^. When diploid cells are starved, Ume6 and Ime1 interact, which drives the transcription of the EMGs ^13,14^. The kinase Rim11 and GSK-3β homolog also plays critical role in the activation of EMGs. Rim11 phosphorylates both Ime1 and Ume6, which is required for the two proteins to interact and to activate the transcription of EMGs^16–18^. Moreover, Rim11 directed phosphorylation of Ime1 is also essential for the Ime1 transcriptional activation domain function when part of the Ume6-Ime1 complex^9,16^. Rim11 can also directly interact with Ime1 *in vitro*^17^. Thus, the Ime1-Ume6-Rim11 regulon drives entry into meiosis. While *IME1* transcriptional control by external and cell-intrinsic cues is well understood and a key determinant for meiotic entry, little is known on how Rim11 is regulated.

Here we show that Rim11 acts as an integrator of multiple signals to robustly regulate EMG transcription and thus meiosis. We find that Rim11 expression and localization is highly regulated in nutrient dependent manner. In nutrient rich conditions Rim11 expression is kept low by PKA and is kept mostly cytoplasmic by TORC1. During starvation Rim11 expression is increased and localized to the nucleus. Importantly, Rim11 nuclear localization is essential for EMG activation and entry into meiosis, but not sufficient. Rim11 requires the presence of Ime1, which acts as the scaffold protein for Rim11 to phosphorylate Ume6. The Rim11 directed Ume6-Ime1 complexes drive the recruitment of coactivators, chromatin remodellers, and basal transcription factors. Single cell analysis and mathematical modelling demonstrate that Rim11 nuclear accumulation can be limiting for EMG induction. Together, our data show that Rim11 acts a as an integrator for nutrient and cell type-specific signals (TORC1/PKA/Ime1) that determine the temporal control of Ume6 phosphorylation, EMG transcription and thereby regulating entry into meiosis.

## Results

### Rim11 phosphorylates Ume6 in diploid cells induced to enter meiosis

Ime1, Ume6, and Rim11 regulate the transcription induction of EMGs ^4,9^. While mating-type and nutrient control of the *IME1* promoter and thus Ime1 expression are well understood, much less is known about how Rim11 is regulated^4^. To obtain insight into Rim11 regulation, we measured Rim11 dependent phosphorylation by western blotting of Ume6, in wild-type *MAT*a/α diploid cells and in *rim11*Δ *MAT*a/α diploid cells as the control. In cells grown in the exponential growth phase (exp) under rich medium conditions we observed no difference in Ume6 migration pattern between WT and *rim11*Δ cells (Figure 1A). When cells were induced to enter meiosis in sporulation medium (SPO) using a synchronous meiosis protocol, we detected a slower Rim11-dependent migrating Ume6 form at 2, 4, 6 and 8 hours in SPO peaking at 4 hours in SPO, but no difference was observed at 0, 1 and 24 hours in SPO (Figure 1A). Our results are consistent with previous work that showed Ume6 is phosphorylated in a nutrient dependent manner^16^. We conclude that Rim11 directed phosphorylation of Ume6 occurs at specific time points during meiosis, suggesting that Rim11 is regulated by specific meiotic entry signals.

**Figure 1.**
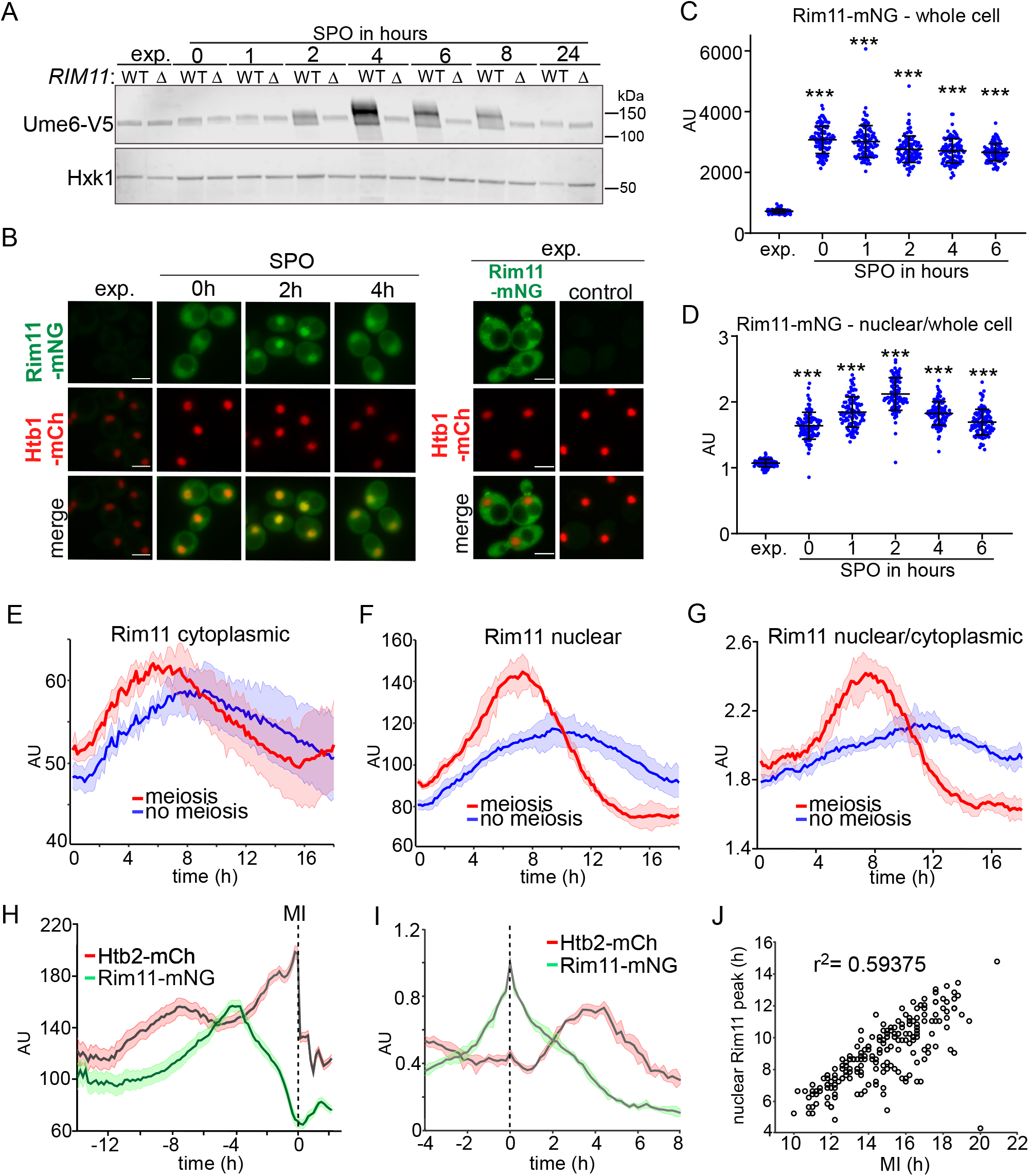
Rim11 expression and localization dynamics prior and during meiosis. (**A**) Western blot of Ume6 in wild-type and *rim11*Δ. Cells harbouring Ume6 epitope tagged with V5 in WT or *rim11*Δ (FW1208 and FW10033) were grown untill exponential phase (exp.) or were grown to be induced to enter meiosis in sporulation medium using a protocol where most cells underwent meiosis (typically 80% or more). (**B**) Localization of Rim11 in exponential growth (exp.) and in cells entering meiosis. We strain was used epitope tagged Rim11 with mNeongreen (Rim11-mNG) and histone H2B epitope tagged with mCherry (Htb1-mCh) (FW10297). A negative control strain harbouring Htb2-mCh only was included (FW7794). **(C)** Quantification of whole cell Rim11-mNG intensity at the different time points. (**D**) Similar as C, except that nuclear over whole cell concentration are displayed. (**E-G**) Quantification of intensity of Rim11 using live cell imaging set up using the strain described in B. Indicated are the mean traces of cells that underwent meiosis and cells that did not underwent meiosis (non-meiotic). Shown are the Rim11 cytoplasmic concentrations (E), nuclear (F), and nuclear/cytoplasmic (G). (**H**) Same data of Rim11 and Htb1 intensities were aligned according the first meiotic division (MI). **(I)** Quantification of Rim11 and Htb1 intensity. Cells were aligned according to the Rim11 peak concentration. (**J**) Scatter plot displaying the timing of Rim11 nuclear peak versus the timing of MI divisions.

### Rim11 expression and localization differ greatly between mitotic and meiotic cells

Given that the Rim11 substrate Ume6 is a DNA binding protein that associates with specific DNA sequence motifs and exclusively resides in the nucleus, we examined whether and how Rim11 nuclear levels are regulated ^19,20^. We fused Rim11 to mNeongreen (Rim11-mNG) and determined its expression and localization in nutrient rich conditions and in cells entering meiosis. To determine the nuclear signal of Rim11 we also expressed histone H2B signal fused to mCherry (Htb1-mCh). Rim11 levels markedly increased in cells entering meiosis compared to cells grown in nutrient rich conditions (exp) (Figure 1C and Figure S1A). This increase in meiotic cells was also reflected at the *RIM11* mRNA levels suggesting that *RIM11* transcription is regulated in a nutrient dependent manner (Figure S1B). Additionally, we observed notable changes in Rim11 localization. In cells grown in nutrient rich conditions (exp.) the majority of Rim11 resided in the cytoplasm, while cells entering meiosis showed strong nuclear localization of Rim11 (Figure 1D and Figure S1A). Both Rim11 nuclear intensity and nucleocytoplasmic ratio were the strongest at 2 hours in SPO. These data show that Rim11 mRNA/protein levels and localization differ greatly between mitotic and meiotic cells.

To correlate meiosis with Rim11 nuclear levels in the same cell, we tracked single cells using fluorescence time lapse microscopy of Rim11-mNG (Supplementary File 1). Consistent with the time course analysis of Rim11 localization (Figure 1D), we observed that Rim11 expression and nuclear localization increased over time in cells that underwent meiosis (Figure 1E-1G and Figure S1C). Strikingly, we found that cells that underwent meiosis showed significant more Rim11 in the nucleus compared to cells that did not undergo meiosis (Figure 1E-1G). This indicates that increased Rim11 expression and nuclear accumulation is specific for meiotic cells and not a general starvation induced signal.

Computational alignment of meiotic cells to the onset time of the first meiotic division revealed that nuclear Rim11 peaked prior the first meiotic division, after which Rim11 expression and nuclear localization decreased when cells progress further into meiosis (Figure 1H). The meiotic cells experienced a peak in Rim11 nuclear levels between 3 and 6 hours before the onset of meiosis I (MI) (Figure 1I). By aligning meiotic cells to the time when Rim11 nuclear levels peak we also observed that Rim11 peaks prior to the increase in histone H2B signal, which is indicative of premeiotic DNA replication (Figure 1I). In line with the above observations, we found that there is a good correlation between the nuclear Rim11 peak signal and timing of MI approximately 6 hours later, suggesting that Rim11 nuclear peaks levels are predictive of when meiosis takes place (Figure 1J). Thus, the increased nuclear concentrations of Rim11 not only correlates with the timing of Ume6 phosphorylation and EMG transcription, but also with the overall propensity to undergo meiosis.

### Rim11 nuclear accumulation is required for entry into meiosis

Next, we examined whether nuclear accumulation of Rim11 is essential for induction of EMG transcription and progression into meiosis. To disrupt nuclear Rim11 accumulation, we adopted the anchor-away method as previously described ^21^. In short, we fused Rim11 to FRB-GFP and Rpl13A to FKBP12 to generate the Rim11 anchor away strain (*RIM11-AA*) (Figure 2A). Using Rim11-AA, we induced Rim11 depletion from the nucleus in pre-sporulation medium (preSPO) and in SPO (*RIM11-AA* + rapamycin). We found that Rim11 was efficiently depleted from the nucleus in preSPO and SPO (Figure 2B, 2C, and Figure S2A). Noteworthy is that the *RIM11-AA* strain showed delayed meiosis only likely because the FRB-GFP tag partially interfered with Rim11 function (Figure S2B). Nuclear depletion of Rim11 (*RIM11-AA* + rapamycin) abolished meiosis, while a large fraction of untreated cells (*RIM11-AA* + mock) completed meiosis (Figure 2D). Rapamycin treatment had no negative effect on Rim11 nuclear localization and meiosis in a strain harbouring Rim11-mNG (Figure 2D and Figure S2C). To assess the effect of the *RIM11-AA* allele on EMG expression, we analysed its impact on the key EMG, the protein kinase Ime2. *IME2* expression was strongly reduced in cells depleted for nuclear Rim11 (*RIM11-AA* + rapamycin) compared to control cells (*RIM11-AA* + mock) (Figure 2E). Consistent with the delay in meiosis observed in the *RIM11-AA* strain, *IME2* levels in control cells (*RIM11-AA* + mock) were lower compared to WT cells at 2 and 4 hours in SPO, however after 48 hours in SPO *IME2* levels in control cells were comparable to WT 4 hours SPO (Figure S2D and S2E). As expected, treatment with rapamycin had no negative effect on *IME2* expression in cell harbouring *RIM11*-mNG instead of *RIM11-AA* (Figure S2D). Importantly, we found that Ume6 phosphorylation was compromised upon Rim11 depletion from the nucleus (*RIM11-AA* + rapamycin). Specifically, after 48 hours in SPO we observed a higher migrating form of Ume6 in the control (*RIM11-AA* + mock) but not in cells with nuclear depletion of Rim11 (*RIM11-AA* + rapamycin) (Figure 2F). We conclude that nuclear accumulation of Rim11 is essential for Ume6 phosphorylation and consequently for the induction of *IME2* transcription and for meiosis.

**Figure 2.**
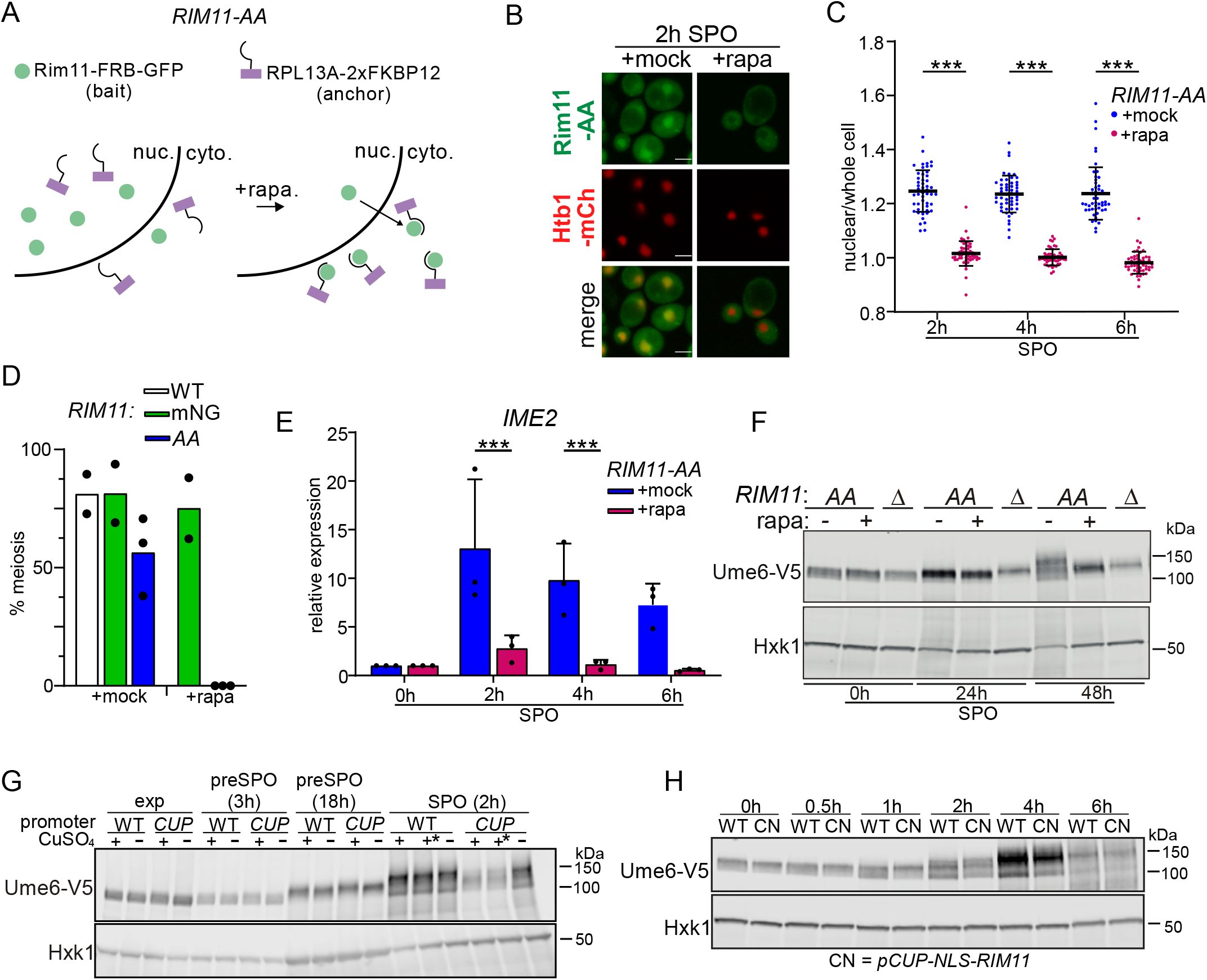
Rim11 nuclear localization is required for meiosis, but not sufficient. **(A)** Schematic of anchoring away of Rim11 from the nucleus. A strain was used harbouring *RIM11* tagged with FRB-GFP, *RPL13A* tagged with 2xFKBP12, gene deletion of *FPR1* (FW11124). **(B)** Rim11 localization in cells that were induced to enter meiosis and that were mock treated or treated with rapamycin for 2 hours. **(C)** Quantification of nuclear over whole cell signal of cells and conditions described in A and B. **(D)** Quantification of cells that underwent meiosis of strain and conditions described in A and B (FW1511, FW11126, FW11124). Meiosis was quantified using DAPI staining. Cells that contained two or more DAPI masses were considered meiosis. The mean of n=2 experiments is shown. At least 200 cells were used for the analysis. **(E)** *IME2* expression determined by RT-qPCR in strain and condition described in A and B. Cells were mock treated or rapamycin treated, and samples were taken at the indicated time points. RT-PCR was performed. Samples were normalized to the time point 0 hours. n=3 biological repeats were performed. The *IME2* signals were normalized over *ACT1*. **(F)** Western blot of Ume6-V5 in cell harbouring the Rim11-AA (FW11371). Cells were induced to enter meiosis and mock treated or rapamycin treated. Samples were taken at the indicated time points for western blotting. Membranes were probed with anti-V5 antibodies. As a loading control Hxk1 was used. **(G)** The effect of overexpression of Rim11 on Ume6 phosphorylation as determined by western blotting. Cells harbouring the *CUP1* promoter (FW10923) and WT *RIM11* promoter (FW11394) and Ume6-V5. Cells were grown and induced to enter meiosis. **(H)** Similar analysis as G except that NLS sequences was fused to Rim11 and expressed from the *CUP1* promoter (FW11394). As a control the WT *RIM11* locus was used (FW11395).

### High Rim11 nuclear levels are not sufficient for meiotic entry

Next, we examined whether Rim11 expression and nuclear localization can drive Ume6 phosphorylation. We induced Rim11 from the *CUP1* promoter (*pCUP1-RIM11*) for 2 hours in various growth conditions. Rim11 was efficiently induced and expressed across whole cells including the nucleus and much higher than the wild-type control (Figure S2G). Despite much higher Rim11 levels in *pCUP1-RIM11* cells, no higher migrating form of Ume6 was detected in cells grown in rich medium (exp.), and in preSPO. After 2 hours in SPO both *pCUP1-RIM11* and WT displayed a higher migrating form of Ume6, but overall Ume6 levels seemed decreased in *pCUP1-RIM11* compared to WT (Figure 2G). We further forced nuclear Rim11 by fusing nuclear localization sequence (NLS) to the amino terminus of Rim11 expressed from the *CUP1* promoter (*pCUP1-NLS-RIM11*) (Figure S2H). *pCUP1-NLS-RIM11* induced cells showed a comparable Ume6 migration pattern to WT, and the onset meiosis was somewhat delayed in *pCUP1-NLS-RIM11* cells (Figure 2H and Figure S2I). Thus, increasing Rim11 levels or forcing Rim11 to nucleus is not sufficient to induce an earlier onset of meiosis suggesting that additional signals are required.

### Rim11 expression and localization are regulated by TOR and PKA signalling

Having established that Rim11 localization to the nucleus is required (yet not sufficient) for meiosis, we next examined how Rim11 localization is regulated. We first determined how Rim11 localization is affected in different nutrient environments. We shifted cells grown in rich medium (YPD) to a rich medium lacking a carbon source (YP), or from preSPO medium to either SPO, SPO plus glucose (Figure 3A and 3B). Rim11 nuclear localization was significantly increased when cells were shifted from YPD to YP. Conversely, nuclear Rim11 intensity was significantly reduced when cells were shifted to SPO + glucose instead of SPO.

**Figure 3.**
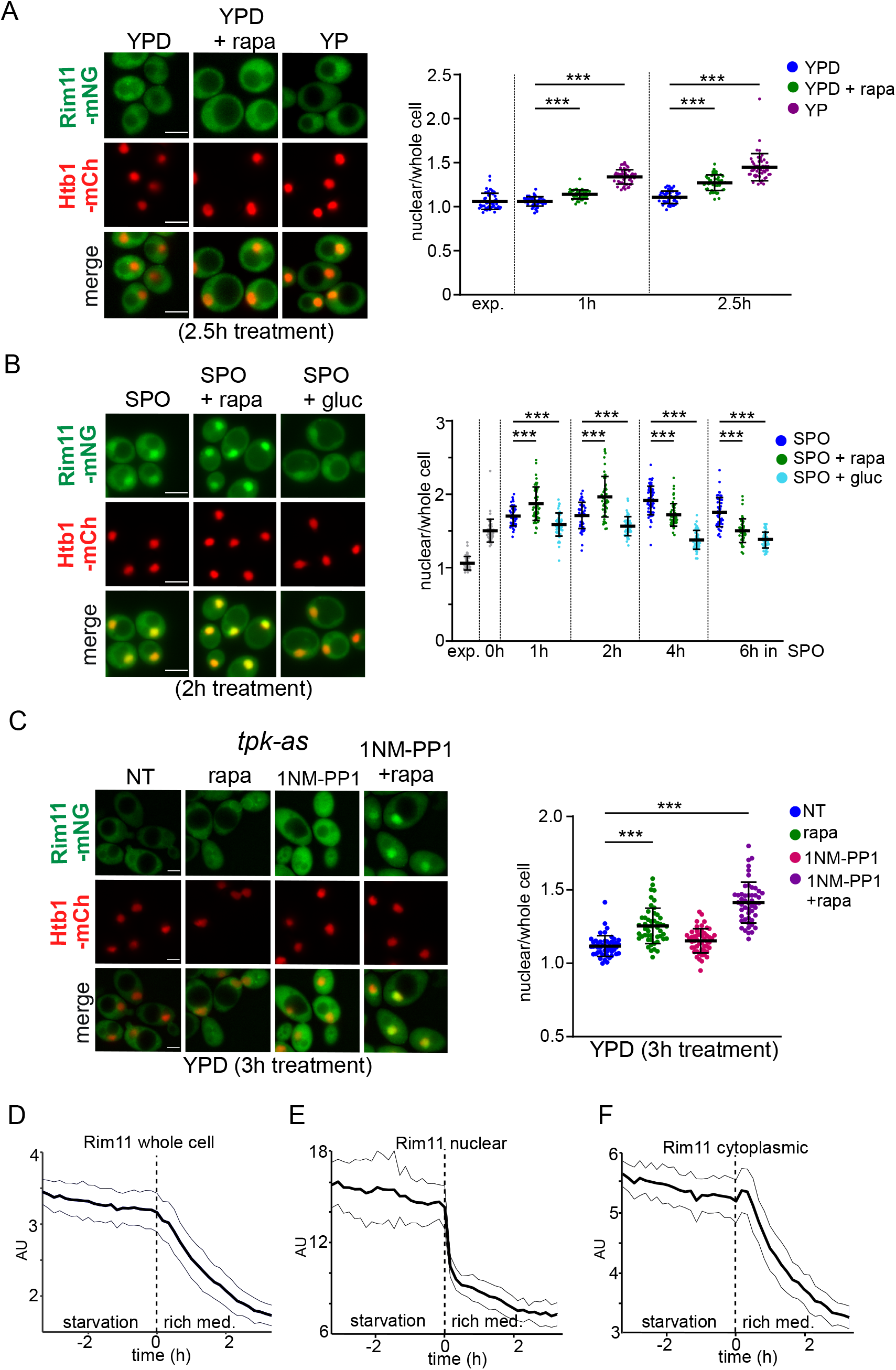
TORC1 and PKA control Rim11 localization and expression. **(A)** Rim11-mNG (FW10297) localization upon rapamycin treatment. Cells were grown in rich medium to exponential growth (exp.) treated with rapamycin for the indicated times. Representative images are shown (left) and nuclear over whole cell quantification is shown (right). **(B)** Similar analysis as in A, except that cells were induced to enter meiosis in SPO, SPO + rapamycin, and SPO + glucose. **(C)** Similar analysis as in A, except that Rim11-mNG also contained *tpk1-as* allele described previously (FW10483) was used for the analysis. Cells were grown in rich medium to exponential growth (exp.) treated with NMPP1, rapamycin or both compounds for three hours. (**D-F)** Rim11 localization dynamics during shift from starvation to rich medium in whole cell, nucleus, and cytoplasm (FW10297). Shown are the Rim11-mNG signal. The data were centred on the starvation to rich medium transition. The mean signals and confidence interval are indicated.

The major nutrient sensing kinase complex and growth regulator in yeast cells is target of rapamycin complex 1 (TORC1). We determined whether TORC1 mediates Rim11 localization by inhibiting TORC1 with rapamycin treatment in YPD and SPO. Cells treated with rapamycin showed increase Rim11 localization compared to nontreated cells (Figure 3A). Likewise, cells shifted to SPO plus rapamycin showed increased nuclear localization of Rim11, and more rapid increase in Ume6 phosphorylation (Figure 3B and Figure S3A). We conclude that a fermentable carbon source (glucose) in the growth medium and TORC1 are important for retaining Rim11 outside the nucleus.

Besides TORC1, PKA is also a key signalling kinase complex for nutrient sensing (such as glucose) in eukaryotes including yeasts^22^. Both PKA and TORC1 signalling play also a central role in regulating the decision to enter meiosis^8,23,24^. When both PKA and TORC1 signalling are inhibited, diploid cells induce transcription from the *IME1* promoter allowing cells to activate EMG transcription and to enter meiosis. Our data suggest that the nutrient signals that regulate the *IME1* promoter also control Rim11 localization (Figure 3A and 3B). We next examined how PKA controls Rim11 localization and expression. We inhibited PKA activity with an ATP analogue 1NM-PP1 in a strain carrying the analogue sensitive *tpk1-as* allele and gene deletions in *TPK2*, and *TPK3*. When we inhibited PKA in rich medium conditions (YPD), Rim11-mNG overall signal increased, while Rim11 nuclear localization remained unchanged (Figure 3C and Figure S3B). Thus, PKA regulates the overall expression levels of Rim11 but not its localization. Similar as we observed in Figure 3B, Rim11 nuclear signal increased upon inhibition of TORC1 with rapamycin, while Rim11 expression only marginally increased (Figure 3C and Figure S3B). When both PKA and TORC1 were inhibited, we observed a significant increase in Rim11 expression and nuclear localization (Figure 3C and Figure S3B). We conclude that PKA predominantly regulates Rim11 expression, while TORC1 has a prime role in controlling Rim11 localization in cells.

### PKA and TORC1 represses Rim11 mRNA and protein accumulation

How do PKA and TORC1 control Rim11 expression? We first examined how *RIM11* mRNA levels are regulated. We assessed the *RIM11* expression pattern from a published dataset that determined expression profiles upon repletion from glycerol rich medium to glucose rich medium with either PKA inhibited using the PKA^AS^ allele, TORC1 inhibited, or both signalling pathways inhibited (Figure S4A)^25^. The shift to glucose medium resulted in a rapid reduction in *RIM11* mRNA levels. However, in PKA or TORC1 inhibited (1NM-PP1 or rapamycin treatment) cells *RIM11* mRNA increased. Additionally, *RIM11* mRNA further increased when both pathways were inhibited indicating a synergistic effect. As expected, *ACT1* levels did not differ between the different treatments. We conclude that PKA and TORC1 repress *RIM11* mRNA expression.

To confirm these results at the single cell level, we analysed the dynamics of Rim11 localization in response to nutrient availability using live cell imaging (Figure 3D, 3E, and 3F). After starvation onset, we observed that nuclear Rim11 is progressively accumulated over a period of at least 7 hours before its levels plateaued and slowly declined (Figure 1F). During the transition from starvation to nutrient rich conditions, Rim11 was downregulated gradually (50% reduction in 70 +/− 40 minutes) (Figure 3D). Interestingly, Rim11 depletion was much faster in nucleus than in the cytoplasm (18 ± 6 min minutes for a 50% reduction in nuclear levels vs 72 ± 40 min minutes for 50% reduction in cytoplasmic levels) (Figure 3E and 3F), suggesting a switch-like change in the nuclear to cytoplasmic ratios. This difference in the regulation of the nuclear and cytoplasmic pools of Rim11, further supports the idea that nuclear Rim11 levels are tightly regulated through different pathways, TORC1 and PKA, respectively.

Previous work suggested that PKA also directly phosphorylates amino-terminal residues in Rim11 on serine 5, 8 and 12. A mutant of Rim11 with the three serines substituted with alanines (*RIM11^3SA^*) displayed increased Rim11 kinase activity in glucose medium, which mis-regulates the onset of meiosis. We examined whether the *RIM11^3SA^* affected Rim11 localization, expression, and the onset of meiosis (Figure 4A, 4B and Figure S4B). We found that Rim11 protein expression was increased in the *RIM11^3SA^* mutant, but localization pattern during entry into meiosis was not affected in a synchronous meiosis (Figure 4A, 4B, and Figure S4B). Also, we performed live cell imaging with the *RIM11^3SA^* mutant. Consistent with the still images, we found that whole cell intensity before MI, but not nuclear intensity, was higher for *RIM11^3SA^* compared to *RIM11*^WT^ (Figure S4C). Interestingly, we noted that the onset of meiosis was more rapid using live cell imaging set up suggesting that in this setup the *RIM11^3SA^* mutant mis-regulated the timing of meiosis (Figure S4D). Both Rim11 and *RIM11^3SA^* cells displayed reduced nuclear localization in SPO with glucose (Figure 4B and S4E). We conclude that PKA mediated phosphorylation of Rim11 is likely destabilizes Rim11 and reduces its protein levels and propensity to undergo meiosis but had little effect on Rim11 nuclear localization.

**Figure 4.**
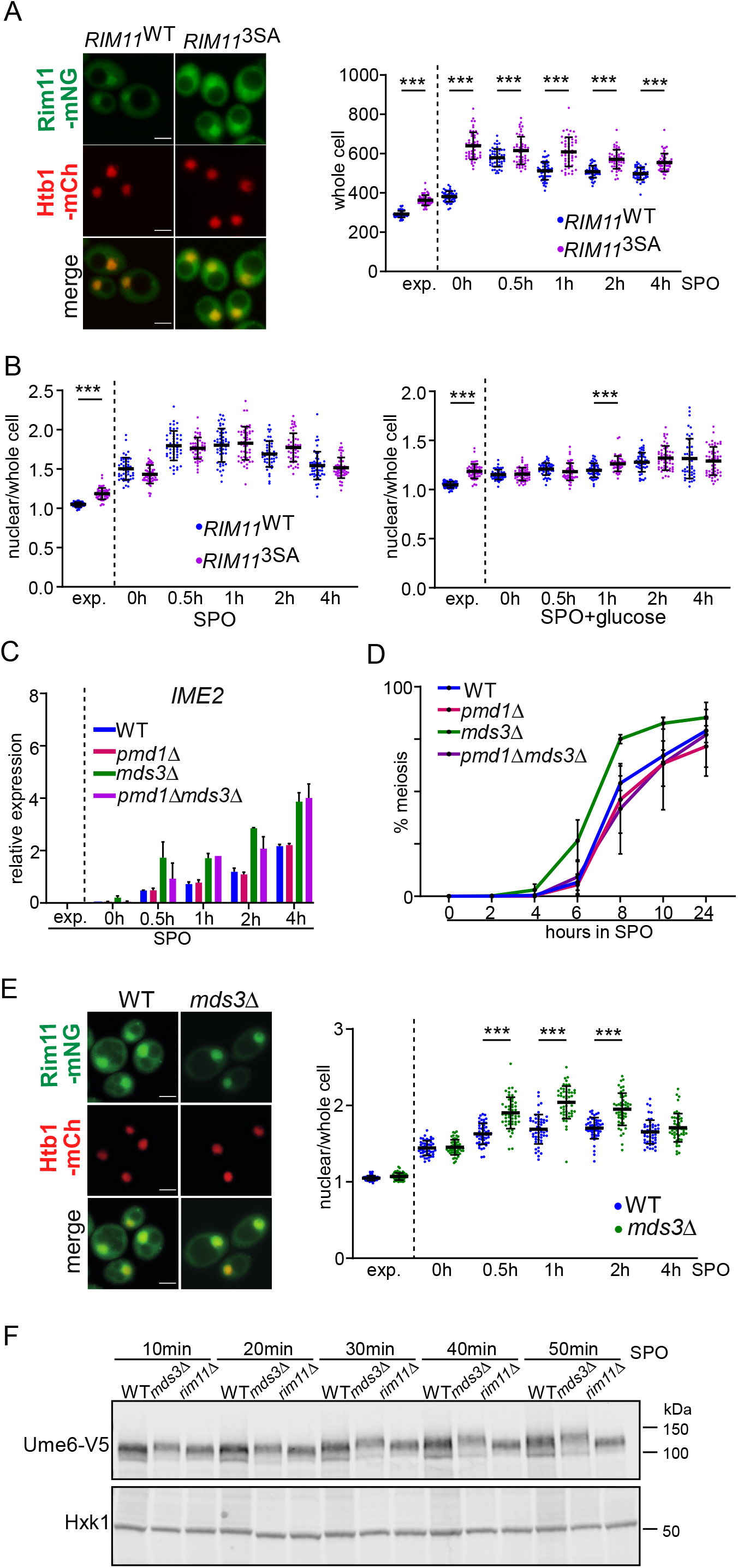
PKA and TORC1 control Rim11 via distinct mechanisms. **(A)** Localization of Rim11^WT^ and Rim11^3SA^ (FW10776 and FW10778). Cells were induced to enter meiosis and samples were taken at the indicated time points, whole cell signal of Rim11-mNG was determined. Shown are example images for the 0 hour time points (left), and the whole cell quantification (right). **(B)** Similar analysis as in A but showing nuclear over whole cell signal of cells induced to enter meiosis in SPO (left) or SPO plus 2% glucose (right). **(C)** *IME2* expression in cell induced to enter meiosis in WT, *pmd1*Δ, *mds3*Δ, and *pmd1*Δ*mds3*Δ. (FW10297, FW10716, FW10718, FW10762). *IME2* expression signals were normalized over *ACT1*. The mean signals of n=3 repeats are shown. **(D)** Onset of meiosis of strains described in C. The mean signals of n=3 repeats are shown. **(E)** Rim11-mNG nuclear over whole cell localization in WT and *mds3*Δ cells induced to enter meiosis (FW10297, FW10718). **(F)** Ume6 migration as determined by western blotting in WT, *mds3*Δ, and *rim11*Δ (FW1208, FW11251, FW10033). Cells were induced to enter meiosis and samples were taken at the indicated time points. Membranes were probed with anti-V5 antibodies and Hxk1 antibodies as a loading control.

### Mds3 promotes the Rim11 localization via TORC1

Next, we turned to analysing how TORC1 regulates Rim11 nuclear localisation. TORC1 is a multi-subunit complex with several downstream effectors that could be involved in mediating TORC1 signalling to Rim11^26^. We identified the paralogs Mds3 and Pmd1 as two potential candidates to regulate Rim11 and meiosis. *mds3*Δ and *pmd1*Δ supress the meiotic defect of *mck1*Δ cells, another GSK-3 homologue that can phosphorylate Ume6 ^27^. Also, *mds3*Δ and *pmd1*Δ cells display an aberrant expression of meiotic genes during vegetative growth. Lastly, Rim11 and Pmd1 potentially interact ^28,29^.

To determine whether Mds3 and Pmd1 are important for meiosis and possibly regulate Rim11 localization, we determined *IME2* expression and meiosis in *mds3*Δ and *pmd1*Δ single and double mutant cells (Figure 4C and 4D). We found that *IME2* expression was more rapidly induced in *mds3*Δ cells, but not in *pmd1*Δ or *mds3*Δ *pmd1*Δ cells. Also, the onset of meiosis was more rapid in *mds3*Δ cells, but not in *pmd1*Δ or *mds3*Δ *pmd1*Δ cells (Figure 4D). These data suggest Mds3, but not Pmd1, represses *IME2* expression, and consequently the onset of meiosis.

Next, we examined whether Mds3 controls Rim11 localization and Ume6 phosphorylation. Consistent with the *IME2* expression data, Rim11 localized to nucleus more in *mds3*Δ compared to the control for 0.5, 1, 2, 4 hours in SPO time points (Figures 4E and S4F). We noted a slightly higher migrating form of Ume6 protein at 0.5 hours in SPO in *mds3*Δ compared to the control cells, but not at other time points (Figure S4G). A fine time course revealed that slower Ume6 migration forms were consistently more abundant in *mds3*Δ cells compared to WT in the first hour after shifting cells to SPO (Figures 4F). It is worth noting that *mds3*Δ cells still displayed Rim11 nuclear localization changes upon starvation, suggesting that additional molecular mechanisms are important for controlling Rim11 localization.

Mds3 has been implicated in TORC1 mediated signalling ^30^. To determine whether Mds3 acts indeed via TORC1, we treated cells entering meiosis with rapamycin and determined *IME2* expression (Figure S4H). Consistent with the observation that the rapamycin treatment led to the faster Rim11 localization to the nucleus, we found that *IME2* expression was more rapidly induced (Figure 4C and S4H). In *mds3*Δ cells, however, we observed no difference in the induction of *IME2* expression between rapamycin and mock treated cells. We conclude that although Mds3 is not essential for meiosis entry, it specifically promotes signalling from TORC1 to Rim11 to control the timing of nuclear Rim11 localization and EMG expression.

### Rim11 kinase activity and Ime1 contribute to Rim11 nuclear accumulation

Rim11 phosphorylation of Ime1 and Ume6 has been suggested to have multiple functions. First, Rim11 phosphorylation of Ime1 and Ume6 promotes the formation of Ume6-Ime1 complexes, which is critical for driving transcriptional activation of EMGs^16,17^. Second, Rim11 phosphorylation of Ume6 is thought to relieve Sin3-Rpd3L co-repressor complex from Ume6^9,18^. Third, Rim11 phosphorylation of Ime1 is required for the transcription activation domain function of Ime1 which drive induction of EMG transcription^17^. Given that both Rim11 substrates (Ime1 and Ume6) act in the nucleus, we addressed whether Rim11 kinase activity and its interaction with Ime1 contribute to Rim11 localization pattern and Rim11’s ability to phosphorylate Ume6.

To examine whether Rim11 kinase activity is important for its localization, we mutated its autophosphorylation site Y199 to F (*RIM11^Y^*^199^*^F^*), which impairs Rim11 kinase activity. Indeed, *RIM11^Y^*^199^*^F^*was impaired for Ume6 phosphorylation and induction of *IME2* expression (Figure S5A and S5B). While *RIM11^Y^*^199^*^F^* entered the nucleus, the nuclear over whole cell ratio was significantly reduced compared to the control (Figure 5A). Given that Ime1 is a substrate for Rim11 phosphorylation and can directly interact with Rim11 and Ume6, we also examined the contribution of Ime1 in Rim11 localization. We found that in *ime1*Δ cells Rim11 nuclear localization was also significantly reduced (Figure 5B). We conclude, that Rim11 kinase activity and Ime1 contribute to Rim11 accumulation in the nucleus.

**Figure 5.**
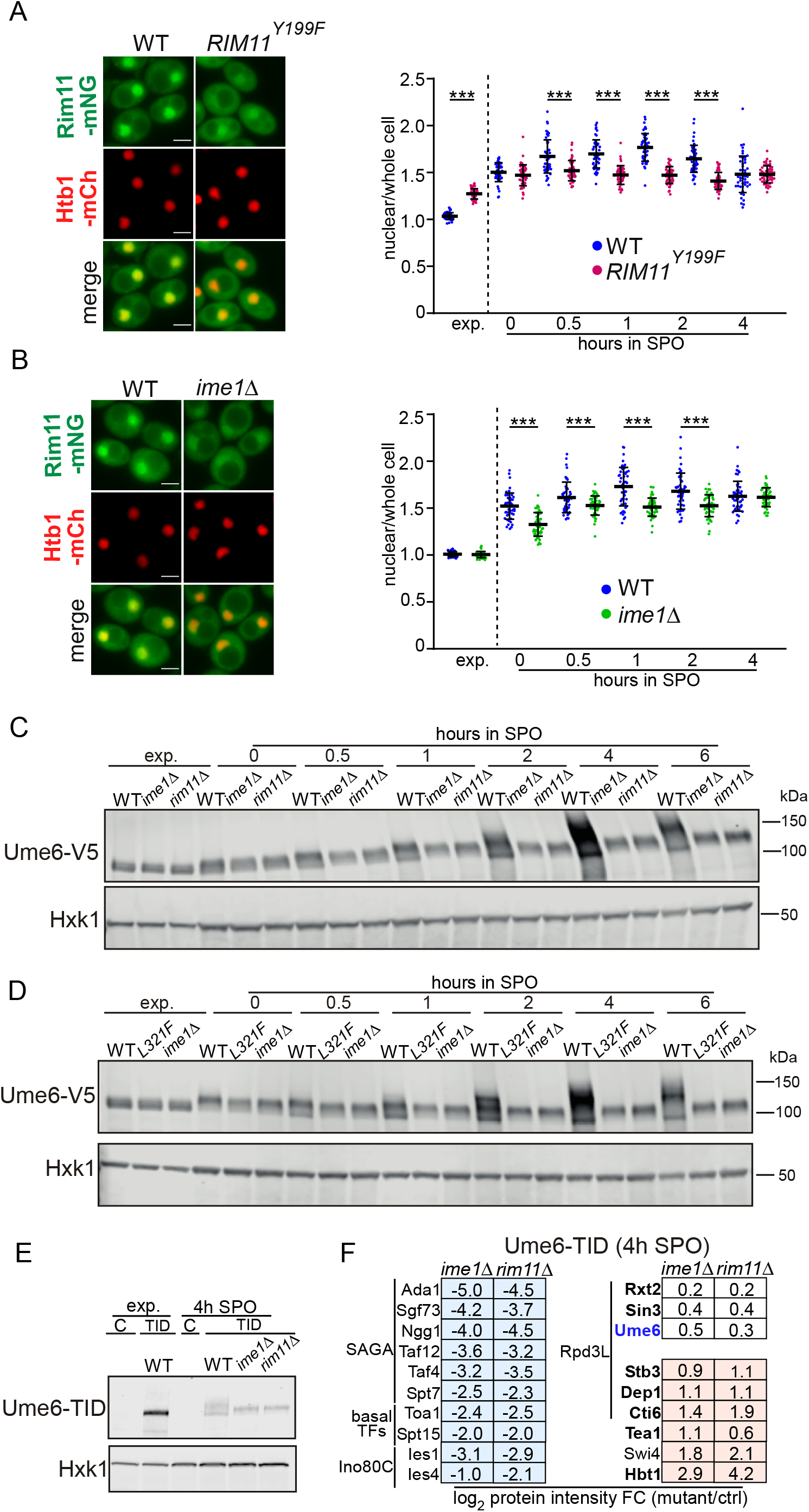
Ime1 is essential for Rim11 directed Ume6 phosphorylation. **(A)** Rim11-mNG nuclear over whole cell localization in Rim11^WT^ and Rim11^Y199F^ (FW10776, FW10983) in cells induced to enter meiosis. Example imagines are shown (left), and quantification (right). **(B)** Same analysis in A but comparing WT and *ime1*Δ cells (FW10776, FW10918). **(C)** Ume6 expression and migration as determined by western blotting in *ime1*Δ and *rim11*Δ cells induced to enter meiosis (FW1208, FW11097, FW10033). Membranes were probes with anti-V5 antibodies and Hxk1 antibodies. **(D)** Similar analysis as C but using the *IME1*^WT^, *IME1*^L321F^ and *ime1*Δ (FW11231, FW11233, FW11097). **(E-F)** TurboID analysis of Ume6. WB showing expression of TurboID tagged Ume6 (Ume6-TID) in exponential phase in WT, and in cells induced to enter meiosis in WT, *ime1*Δ and *rim11*Δ (FW11422, FW11424, FW11426) in E. Log2 protein intensity of *ime1*Δ or *rim11*Δ over control. Listed are proteins involved in transcription or known to interact with Ume6. Significant proteins are identified by permutation-based FDR-corrected *t*-test (threshold: *P* value= 0.05 and S0 = 0.1)

### Ime1 is required for Rim11-dependent Ume6 phosphorylation

Our data showed that Ime1 contributes to the nuclear localization of Rim11. Given that Ime1 can directly interact with both Ume6 and Rim11, we explored whether Ime1 also has a role in Ume6 phosphorylation^11,12,17^. Strikingly, our data showed that Ime1 can be phosphorylated by Rim11 as soon it is expressed, suggesting Ime1-Rim11 interaction precedes Ume6 phosphorylation. When we induced Ime1 from the *CUP1* promoter (*pCUP1-HA-IME1*) in *RIM11*^WT^ and *rim11*Δ cells, in both pre-SPO and SPO conditions a large fraction of Ime1 protein migrated higher in a Rim11 dependent manner (Figure S5C and S5D). Cells expressing Ime1 under nutrient rich conditions (exp.) did not display a difference in the Ime1 migration patterns between *RIM11*^WT^ and *rim11*Δ (Figure S5C).

Given that the Rim11-dependent phosphorylation of Ime1 precedes Ume6 phosphorylation, we determined whether Ime1 anchors Rim11 for Ume6 phosphorylation. Strikingly, starvation exposed *ime1*Δ cells displayed no Rim11 dependent Ume6 migration shift, indicating that Ime1 is required for Rim11 phosphorylation of Ume6 (Figure 5C). To dissect whether the Ime1 interaction with Rim11 is required for Ume6 phosphorylation, we used a previously characterized *IME1* missense mutant (*IME1^L^*^321^*^F^*) which impairs the interaction with Rim11. Like *ime1*Δ cells, *IME1^L^*^321^*^F^* cells were also defective in Ume6 phosphorylation and displayed no induction of *IME2* transcription (Figure 5D and S5E). We conclude that Rim11 dependent phosphorylation of Ume6 requires Ime1. Ime1 likely acts as an anchor or scaffold protein for Rim11. In addition to TORC1 and PKA signals, our data show that Ime1 is a third signal requirement for Rim11 dependent Ume6 phosphorylation and EMG transcription. This surprising result suggest that Rim11 is the central signal integrator for Ume6 phosphorylation and EMG transcription.

### TurboID of Ume6 reveals nutrient dependent interactions with co-repressors and co-activators

To gain insight into mechanism by which Ime1-Rim11-Ume6 activate EMG transcription, we used proximity labelling of Ume6 to map changes in the local transcription and chromatin environment. Specifically, we tagged Ume6 at the carboxy terminus with TurboID for proximity biotin labelling (Ume6-TID). In short, we grew cells in presence of biotin, generated denatured protein extracts from which we purified biotinylated proteins with streptavidin beads. Subsequently, the associated proteins were analysed by MS using label free quantification. The Ume6-TID showed a Rim11 and Ime1 dependent phosphorylation shift demonstrating that the TurboID tag on Ume6 did not affect phosphorylation (Figure 5E). We first generated Ume6-TID profiles by isolating the biotinylated proteins from nutrient rich conditions and from cells entering meiosis (4h SPO). For the analyses we selected proteins that were significantly enriched in the Ume6-TID compared to the untagged control and that are known to associate with Ume6 or are involved in regulating chromatin and transcription. As expected, cells entering meiosis displayed a signature of proteins corresponding to activation of transcription (Figure S5F and Table S1). Six subunits of the coactivator SAGA (e.g. Spt7, Ada1), two subunits of the chromatin remodeller Ino80C (Ies1, Ies4), several basal transcription factors (Spt15, Toa1) and transcription elongation factors (FACT, Spt5, Spt6) were significantly enriched. Conversely, Tod6 subunit of Sin3-Rpd3L which is known to interact with Ume6, was more enriched in the Ume6-TID sample from nutrient rich conditions, while Sin3 and Rxt2 of the Sin3-RpdL co-repressor were not significantly changed. Tea1, Ty1 enhancer activator I and known interactor of Ume6, was strongly enriched in nutrient rich conditions. Noteworthy, Rim11 and Ime1 were not detected, likely because Ime1 associates with amino-terminus region Ume6 while the TurboID was added to the carboxy-terminus of Ume6. We conclude that Ume6 proximity labelling can detect nutrient dependent interactions with protein complexes that repress (Sin3-Rpd3L) or activate transcription (SAGA and Ino80C).

### Rim11 and Ime1 are both required for SAGA recruitment to EMGs

Next, we compared profiles of proteins labelled by Ume6-TID between WT, *ime1*Δ and *rim11*Δ in starved cells (4h SPO). The *ime1*Δ and *rim11*Δ cells showed decreased enrichment for the six subunits of the coactivator SAGA, two subunits of chromatin remodeller Ino80C, and several basal transcription factors (Spt15, Toa1) compared to WT (Figure 5F and Table S1). Moreover, three subunits of Sin3-Rpd3L (Cti6, Dep1 and Stb3) were enriched in the *ime1*Δ and *rim11*Δ cells compared to the WT, while Sin3 and Rxt2 of Sin3-Rpd3L were not significantly changed, supporting the idea that Ume6 phosphorylation affects the interactions with Sin3-Rpd3L. Based on the Ume6 proximity labelling analyses, we conclude that both Ime1 and Rim11 are required for partially disassembly of Sin3-Rpd3L, and recruitment of SAGA co-activator and basal transcription factors to the promoters of EMGs.

### Rim11 and Ime1 can be dispensable for EMG induction and meiosis

The Rim11 phosphorylated forms of Ume6 and Ime1 make up the transcription activator complex for EMG induction and initiation of meiosis^17^. It is not known whether Rim11 and Ime1 act exclusively in EMG activation within the Ume6 regulon. Indeed, Rim11 has been implicated to regulate lipid synthesis and replication stress ^31,32^. We next determined whether Rim11 and Ime1 requirement for meiosis is dispensable by rewiring the Ume6 regulon.

We tested whether we could drive EMG expression and meiosis by tethering Ime1 to the *UME6^T99N^* allele in the absence of *RIM11* (Figure 6A and Figure 6B). *UME6^T99N^* allele impairs the interaction with Ime1, impairing EMG transcriptional activation. With the use sfGFP-Ime1 and GFP nanobodies tagged *UME6^T99N^* (*UME6^T99N^*-αGFP), it is possible to tether Ime1 to Ume6^T99N^-αGFP and thereby drive EMG transcription and meiosis as shown previously (Figure 6A and 6B). However, *rim11*Δ cells harbouring sfGFP-*IME1*/*UME6^T99N^*-αGFP displayed no EMG activation and no meiosis, which is consistent with the model that Rim11 phosphorylation of Ime1 turns Ime1 into transcriptional activation domain that is essential for driving EMG transcription and meiosis^17^.

**Figure 6.**
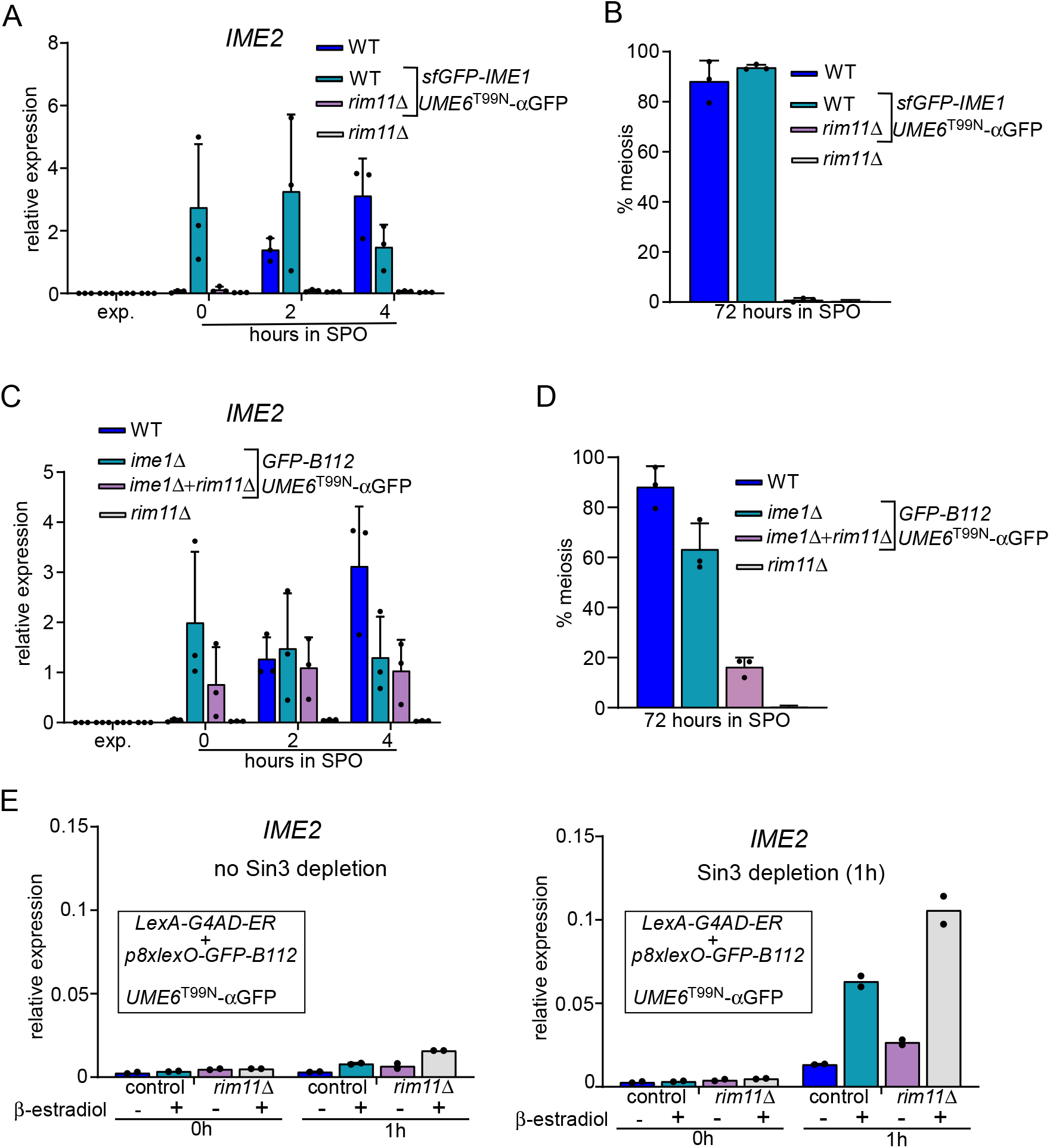
Rewiring of Ume6 regulon makes Rim11 and Ime1 dispensable. **(A)** *IME2* mRNA expression in cells induced to enter meiosis in WT, *rim11*Δ and strains harbouring sfGFP-Ime1 + Ume6^T99N^-aGFP that are WT for Rim11 or *rim11*Δ (FW1511, FW11432, FW10864, FW9277). *IME2* was normalized over *ACT1*, and n=3 biological repeats are shown. **(B)** Same strains as described A, but the percentage of cells that underwent meiosis is plotted. The mean and SEM of n=3 biological repeats are shown. **(C)** Same set up as A, except that sfGFP-Ime1 was replaced with GFP-B112 activation domain (FW1511, FW11414, FW11428, FW9277). **(D)** Same strains as C but plotted are the percentage of cells that underwent meiosis. The mean and SEM of n=3 biological repeats are shown. **(E)** *IME2* mRNA expression in cells grown in exponential growth phase. Similar strains setup as C except that GFP-B112 was under control of LexAG4AD-ER and fused to promoter with eight lexO sites (p8xlexO) (UB36868, UB37219). Additionally, *IME2* mRNA expression was determined in the presence (left) or absence (right) of Sin3 depletion using auxin induced degron (Sin3-AID). Additionally, cells were treated with 3-indoleacetic acid (IAA) and copper sulphate. The mean signal of n=2 biological repeats is shown.

To bypass the requirement for Ime1 phosphorylation by Rim11, we replaced the *IME1* locus with GFP fused *B112* activation domain (*GFP-B112*), which allows for phosphorylation independent activation of transcription. Cells expressing *GFP-B112* and *UME6*^T99N^-*αGFP* were able to activate *IME2* transcription and underwent meiosis (Figure 6C and 6D). Importantly, *IME2* transcription was also induced in *rim11*Δ cells expressing GFP-B112 and *UME6*^T99N^-αGFP, and approximately 20 percent of cells completed meiosis (Figure 6C and 6D). We conclude that Rim11 and Ime1 can be made dispensable for EMG activation and meiosis, suggesting that Rim11 and Ime1 primarily function in the Ume6 regulon.

Rewiring of the Ume6 regulon is sufficient to drive meiosis in the absence of Ime1 and Rim11. We wondered whether nutrient control of EMG transcription can also be bypassed. We expressed *GFP-B112* from a promoter harbouring eight lex operator sequences (*p8xlexO-GFP-B112*) together with the β-estradiol regulated *GAL4-AD-ER* in *UME6*^T99N^-aGFP cells. When we induced *GFP-B112* with β-estradiol during exponential growth in nutrient rich conditions, we noted that *IME2* expression was only marginally increased suggesting the B112 tethered to Ume6 was not sufficient for EMG induction under these conditions (Figure 6E, left panel). Given the role of Sin3-Rpd3L in repressing EMGs, we reasoned that alleviation of this repression depletion of Sin3-Rpd3L is possibly a prerequisite of EMG activation ^15,18^. To achieve this, we introduced generated an auxin inducible degron tagged allele of Sin3 (SIN3-AID) to conditionally deplete Sin3 (Figure 6E, right panel). Depletion of Sin3 or presence of Tir1 ligase in cells did not result in increased expression of *IME2* (Figure 6E, left panel, and Figure S6). Despite nutrient rich conditions, *IME2* expression was strongly induced in *RIM11^WT^* or *rim11*Δ cells expressing GFP-B112 and depleted for Sin3 (Figure 6E, right panel). We conclude that nutrient control of EMG transcription via Ime1 and Rim11 can be bypassed by tethering an activation domain to Ume6 in cells depleted for Sin3-Rpd3L.

### Single cell analysis of the temporal coordination of the Rim11-Ime1-Ume6 regulon

We showed that Rim11 integrates signals from PKA, TORC1 and Ime1 to regulate induction of EMG transcription and entry into meiosis. The *IME1* promoter also integrates signals from TORC1 and PKA, which control its transcription. How does the variability in Rim11 and Ime1 expression affect the timing of meiosis? To address this, we monitored Rim11, Ime1, and Ume6 nuclear intensities in single cells using live cell imaging. We generated a strain with Rim11, Ime1, and Ume6 fluorophore-tagged at the endogenous loci, with mKOκ, sfGFP, and mTFP1, respectively. Additionally, the strain also carried a construct harbouring an NLS-containing mRuby3 expressed from the *IME1* promoter to label nuclear meiotic divisions. As a readout for EMG expression, we used Ume6 itself because Ume6 directly regulates its own promoter and thus, like EMGs, is transcriptionally induced in early meiosis ^13^.

The four-color strain was able to undergo meiosis and form viable spores (92 ± % viability) in the live cell imaging setup (Supplementary File 2). First, we compared cells entering meiosis with cells that did not complete a meiotic division also here referred to as quiescent cells (Figure 6A, and Figure S6A and S6B). As expected, nuclear levels of Rim11 were elevated in meiotic cells compared to non-meiotic cells (Figure S6B). Furthermore, Ime1 and Ume6 nuclear levels were upregulated in cells entering meiosis but not in non-meiotic cells (Figure 6A, and Figure S6A and S6B).

### Rim11 nuclear levels peaks before Ime1 and Ume6

We assessed the nuclear accumulation dynamics of Rim11, Ime1 and Ume6 throughout meiosis. We aligned single cell time series according to the time of the first meiotic division (MI) or according to time of the peak of nuclear Rim11 during early meiotic entry (Figure 7B and 7C). The nuclear accumulation of Rim11 preceded the peak nuclear accumulation of Ime1 and Ume6 regardless of whether cells were aligned to the first meiotic division or the Rim11 peak (Figure 7B). Ime1 levels peaked approximately 3 hours after the Rim11 peak and then plateaued to decline slowly during MI. Interestingly, after reaching its peak, nuclear Rim11 levels declined significantly, reaching their lowest point at the onset of MI. The decline was also observed for Ume6 although with a delay of approximately 3 hours from nuclear Rim11 peak time. We conclude that Rim11 and Ime1 nuclear accumulation starts roughly at the same time, while Rim11 peaks well prior to Ime1 and Ume6. Our data suggest that cells set up the meiotic fate decision by altering Rim11 nuclear levels well prior to the onset of meiosis.

**Figure 7.**
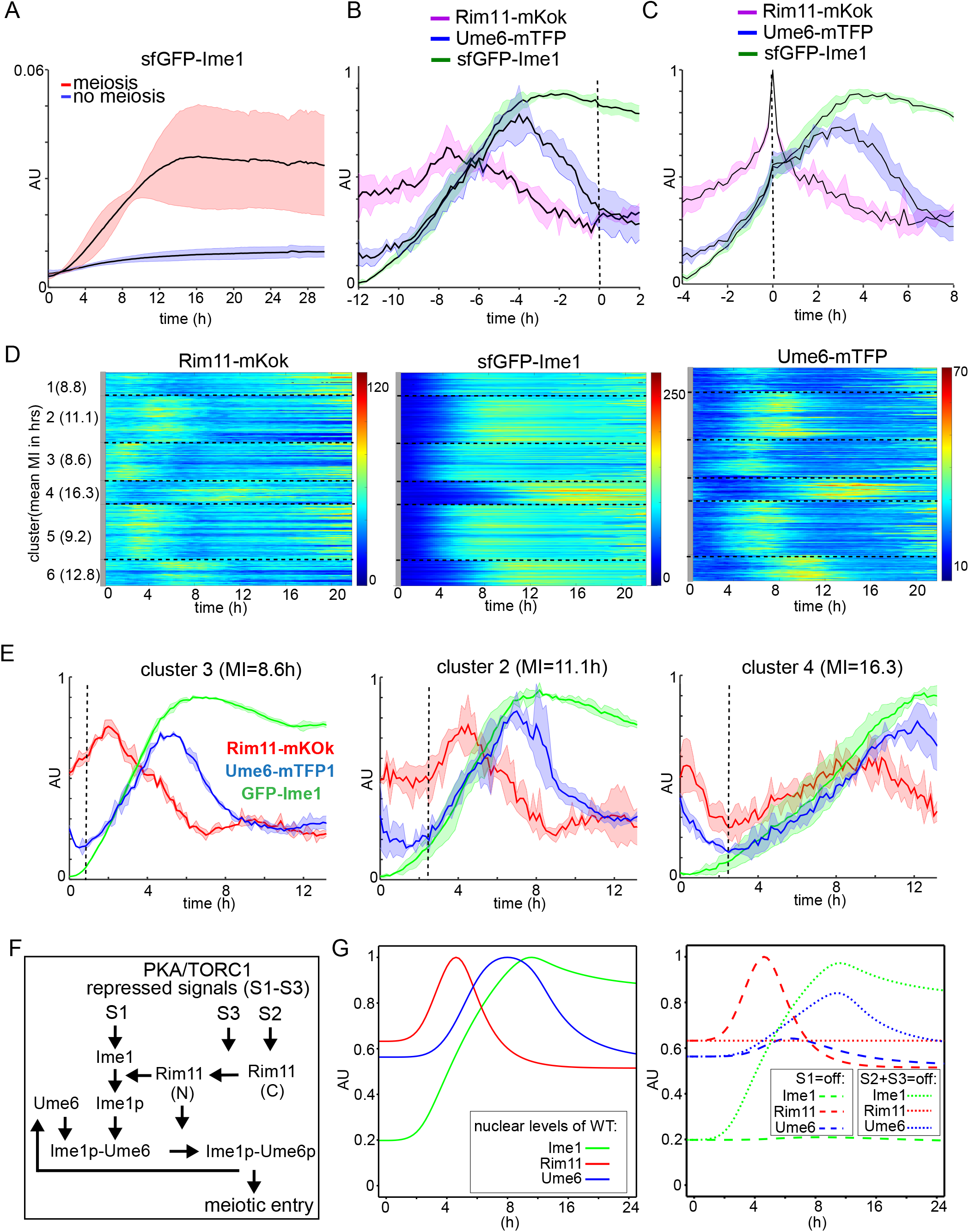
Single cell analysis and modelling reveals the timing dynamics Rim11, Ime1, Ume6 and meiosis. Live cell imaging of the dynamics of Rim11, Ime1, Ume6 in meiotic and non-meiotic cells. We generated a strain with sfGFP-Ime1, Ume6-mTFP, and Rim11-mKok, and pIME1-NLS-mRuby3 (FW11243). **(A)** Nuclear concentrations of sfGFP-Ime1 in meiotic and non-meiotic cells. **(B)** Rim11, Ime1, Ume6 nuclear intensity in a time course aligned according to the MI division. **(C)** Time course data aligned according the Rim11 intensity peak is shown. **(D)** Clustering of single cell traces according to Rim11 nuclear intensity (k=6). **(E)** Mean traces of Rim11, Ime1, Ume6 nuclear intensity for cluster 3 (left), 2 (middle) and 4 (right). **(F)** Scheme of mathematical model. Ime1 expression positively regulated by signal S1, Rim11 expression by S2, and Rim11 nuclear localization by S3. Ime1p and Ume6p represent the Rim11-phosphorylated forms. **(G)** Graph showing the nuclear signal of Ime1, Ume6 and Rim11 from the mean single-cell time series predicted by the model for the wild type (left). Graph showing the mean single-cell time series predicted by the model when switching off the starvation signal to Ime1 (S1) or the starvation signals for Rim11 expression and localisation (S2 and S3) (right).

### Rim11 and Ime1 nuclear levels explain timing variability in meiosis

We used clustering analysis of the single cell time series to determine whether the kinetics in the nuclear Rim11 intensities can explain the heterogeneity in timing observed in meiosis. We clustered (*k*=6) single cell traces for Rim11 and assessed whether the clustering according to Rim11 could reveal different but coordinated patterns for Ime1 and Ume6 accumulation or MI times (Figure 7D). Strikingly, clustering according to Rim11 nuclear levels revealed a consistent pattern in which cells that accumulated Rim11 faster also were advanced in Ime1 and Ume6 accumulation and MI. For example, cluster 2 was the most delayed in Rim11 nuclear accumulation, and so were Ime1 and Ume6 nuclear signals and the onset of MI (Figure 7D and 7E). To confirm the temporal interdependence of Rim11, Ime1, Ume6 peaks with respect to MI (Figure S6C), we calculated the population correlation coefficients and obtained a good correlation between Rim11 nuclear peak and MI (r^2^=0.6), despite the peak of Rim11 occurring approximately 7 hours before MI (Figure S7C, left panel). By contrast the size of meiotic cells was not correlated with MI times (Figure S7D). Both Ime1 and Ume6 peak levels also showed a good correlation with MI (r^2^=0.8 and r^2^=0.83, respectively), which occurred approximately 4 hours before MI (Figure S6C, middle and right panel). The time of Rim11 peak accumulation can largely explain the timing of Ume6 accumulation and meiosis.

### Rim11 nuclear accumulation can be rate-limiting for EMG activation

Previous work suggested that Ime1 is the rate limiting step for induction of EMG transcription and hence meiosis. We wondered whether Rim11 nuclear accumulation can also be rate limiting. As a readout for EMG expression, we followed Ume6 accumulation patterns over time. The different clusters were assessed to determine how Rim11, Ime1, and Ume6 nuclear levels changes correlated (Figure 7D). We noted that in cluster 3, nuclear Rim11 accumulated sharply from the 0-hour time point onwards, while Ume6 and Ime1 (to a lesser extent) were slightly delayed, suggesting that Ime1 was rate limiting in cluster 3 for Ume6 induction (Figure 7E, left panel). In clusters 2 and 5 we noted that Ime1 levels increased well before nuclear Rim11 and Ume6 levels increased, indicating the Rim11 was the rate limiting step in cluster 2 and 5 (Figure 7E, middle panel and Figure S7E). In clusters 4 and 6, we noted that both Rim11 and Ume6 levels initially drop, and subsequently increase at the same time (Figure 7E, right panel; Figure S7E). Ime1 levels, on the other hand, gradually increased over time, suggesting that Rim11 nuclear levels were rate limiting for clusters 4 and 6. Thus, this analysis suggest that Rim11 nuclear concentrations can be rate limiting for EMG induction in some cells, while Ime1 can rate limiting in others.

To determine whether the insights on Rim11 signal integration can explain the effects on EMG expression and meiosis, we constructed a mathematical model that incorporates multi-signal-controlled network including inferred interactions using ordinary differential equations (Figure 7F and Supplementary File 3). We defined three signals repressed by PKA/TORC1, (S1, S2, and S3) representing stimulation of Ime1 expression, Rim11 expression, and Rim11 nuclear localization. The model reproduced the mean single-cell time series for cells with meiosis (Figure 7G, left panel). The model captured the timing of the peaks in nuclear Ime1, Ume6 and Rim11, as well as the amplitude of each signal. The model showed that the signal integration by Rim11 substantially impacts the timing and amplitude of the output signal (Ume6) (Figure 7G, right panel). Removing the starvation signals to Rim11 (S2 and S3) shifted the Ume6 response in time and decreased its expression (Figure 7H). It is worth noting that the model likely overestimated how much “leaky” Rim11 is present when TORC1 and PKA are active. At the single-cell level, when considering the timing of Rim11 and Ime1 nuclear peaks following our single cell data, the model predicted the timing of EMG/Ume6 accumulation as observed in the data (Figure S7F). Overall, the model demonstrated that the signalling network (Rim11-Ime1-Ume6) architecture ensures sensitivity and robustness of EMG transcription, resulting in entry into meiosis.

## Discussion

Despite more than five decades of research, how cells integrate multiple signals to make the decision to enter meiosis and form spores is not well understood. It has been known that GSK-3β ortholog Rim11 is a key part of the regulon for inducing EMG transcription together with Ume6 and Ime1. Here we report the surprising finding that Rim11 integrates multiple signals through distinct mechanisms to control Ume6 phosphorylation and induction of EMG transcription. We found that nutrient availability signals to Rim11 via PKA and TORC1, where PKA controls Rim11 expression levels and TORC1 controls Rim11 nucleocytoplasmic localization. Additionally, Ime1, whose transcription is controlled by nutrient and mating-type signals, acts as the scaffold protein signal for Rim11-dependent phosphorylation of Ume6. Our data suggest that signal integration by Rim11 generates a robust regulatory network structure that controls the timing of EMG transcription, meiosis, and spore formation.

### Rim11 acts as nutrient signal integrator critical for meiotic entry

Our data demonstrate that Rim11 integrates nutrient availability signals via PKA and TORC1 to control its expression and localization, respectively. Inhibiting TORC1 and PKA leads to strong enrichment of Rim11 in the nucleus, which is essential, but not sufficient, for Ume6 phosphorylation and consequently EMG transcription induction. We showed that PKA and TORC1 controls Rim11 expression and localization through distinct mechanisms. This and previous work showed the PKA possibly directly phosphorylates serines at the amino-terminus of Rim11, which likely destabilizes Rim11 protein levels ^33^. Additionally, we also noted that *RIM11* mRNA levels greatly differ between rich medium conditions and starved cells. It was shown that PKA and TORC1 control RIM11 expression levels directly ^25^. Thus, PKA and TORC1 control *RIM11* mRNA accumulation, however, the mechanism by *RIM11* mRNA levels is regulated by nutrient availability remains to be determined.

How does TORC1 retain Rim11 outside the nucleus? We provide evidence that Mds3, in part, retains Rim11 in the cytoplasm. Mds3 is not a well characterized effector protein of TORC1 signalling, however there is some evidence that Mds3 is part of the Sit4 phosphatase branch^30^. Other effectors of TORC1 are likely involved in retaining Rim11 from the nucleus. The Sch9 kinase branch of TORC1 is a plausible candidate since Sch9 is also important for regulating *IME1* transcription and important for entry into meiosis ^8^. However, more work is needed to determine the exact mechanism by which Rim11 localization is controlled by TORC1.

### Multi-layered regulation of Rim11 governs EMG transcription induction

Previous work showed that Ime1 and Rim11 interact directly *in vitro* and Ime1 also interacts with Ume6^17^. Most strikingly, we showed in this work that Ime1 act as the anchor protein for Rim11 to access Ume6 and phosphorylate Ume6. The Ime1 anchor function likely contributes to Rim11 nuclear localization while kinase activity of Rim11 is likely involved in anchoring Rim11 to Ime1. The result is most surprising as previous work suggested that Rim11-depedent phosphorylation can occur in the absence of Ime1, even though the data were not shown^34^.

The scaffold function of Ime1 for Rim11 has various implications for how Rim11-Ime1-Ume6 regulatory network that controls meiotic entry. First, the data show that Ime1 is an integral part of the signal integration cascade for Rim11. The nutrient and mating type control of *IME1* transcription are thus sensed by Rim11 through Ime1. Indeed, expression of Ime1 is not sufficient drive EMG transcription and meiosis in nutrient rich conditions. Our single cell traces show that Rim11 nuclear accumulation can follow Ume6 accumulation, and thus can rate limiting for EMG transcription. The fact that both Rim11 and Ime1 integrate nutrient signals via TORC1 and PKA independently, and yet both are required for Ume6 phosphorylation, generates robustness and specificity in the regulatory network structure controlling EMG transcription.

The single cell data revealed a strong temporal regulation between Rim11, Ime1 and Ume6. Cells accumulate nuclear Rim11 ahead of Ime1 and Ume6. One explanation is that Ime1 phosphorylation occurs first in nucleus, followed by phosphorylation of Ume6 triggering the formation of Ume6-Ime1 complexes. Once Rim11 phosphorylated Ume6-Ime1 complexes are formed, Rim11 exists the nucleus. Exist from the nucleus of Rim11 is also triggered when cells are exposed to nutrient rich conditions, supporting the idea the Rim11-Ime1-Ume6 regulon is rapidly reversible when environmental conditions change.

We constructed a mathematical model of the Rim11, Ime1 and Ume6 regulatory network. This included three input signals to Ime1, Rim11 expression and Rim11 localisation, and the Ime1 scaffolding function in Ume6 phosphorylation. The model captured the data accurately, providing additional support for the proposed signalling network architecture. Modelling also suggests that multi-signal regulation of Rim11 mediates robust cell decision.

### GSK-3 kinases conserved signal integrators

In mammals, glycogen synthase kinase-3β (GSK-3β) has been shown to be crucial for cellular survival, differentiation, and metabolism^35^. The GSK-3 kinases also have been shown to play various roles in meiosis, spermatogenesis, and oocyte development indicating that their roles have been conserved from yeast to humans^12^ ^36–39^. Interestingly, the Ime1 and Ume6 part of the regulatory network is not conserved in mammals.

The localization GSK-3β kinases have been shown to be critical for differentiation of mouse embryonic stem cells (mESC)^40^. Specifically, GSK-3β can translocate to the nucleus to target c-Myc for phosphorylation and degradation. Aberrant nuclear localization of GSK-3β has also been linked to a negative clinical cancer outcome^41^. Like TORC1 and Rim11 yeast, mTor has also been shown to be involved in retaining the localization of GSK-3β to the cytoplasm, and inhibiting mTor leads to nuclear GSK-3β where it activates transcription factors such as c-Myc and Snail^42^. Thus, TORC1/mTor controlled Rim11/ GSK-3β localization, and the targeting of nuclear transcription factors by Rim11/ GSK-3β is likely conserved process from yeast to humans. In contrasts to human GSK-3β, which has over 100 different substrate targets, the role of Rim11 in meiosis is predominantly centred on Ume6 and Ime1 phosphorylation because Rim11 can be made dispensable for meiosis when we rewired the Rim11-Ime1-Ume6 network^35^.

In conclusion, we showed that Rim11 is a master integrator of input signals representing the cell’s metabolic and mating type status (TORC1, PKA and Ime1). The input signals determine whether Rim11 will phosphorylate Ume6 and allowing for Ume6-Ime1 complexes to form which in turn drive EMG transcription and entry to meiosis. Our study provides a completely new view on how the decision into meiosis is regulated, which was thought to be mainly determined by Ime1 expression.

## Supplementary data

**Table S1. Ume6-TurboID dataset**

**Table S2. Genotypes of strains used**

**Table S3. Plasmid used**

**Table S4. Oligo nucleotide sequences**

**Supplementary File 1. Movie of Rim11**

**Supplementary File 2. Movie of Rim11, Ime1, and Ume6**

**Supplementary File 3. Mathematical model**

## Material and methods

### Yeast strain and plasmids

Experiments were carried out with SK1 strain background of *Saccharomyces cerevisiae*. Gene deletions, depletions, anchor away and epitope tagging of strains was done by one-step PCR, and subsequent genetic crosses^43^. Unless stated differently, experiments were performed in liquid cultures at 30°C with 300rpm.

The anchor-away technique was described previously^21^. The SK1 strain with anchor-away modifications was kindly provided by the Adele Marston. The strain contains the ribosomal anchor protein *RPL13A* fused with two copies of the human FKBP12. The target protein *RIM11* was C-terminally tagged with FRB-GFP. Lastly, *FPR1* was deleted allowing the human FKBP12 to bind efficiently to FRB.

For the anchor-away experiment, yeast cultures were grown to induce meiosis using a synchronous meiosis protocol^44^. After 16 hours in BYTA cells were treated with 1μM rapamycin (Sigma-Aldrich, R0395) and incubated for 1h. Then cells were washed and transferred to SPO, OD_600_ =2.5 and treated with 1μM rapamycin.

*UME6*^T99N^ point mutation was generated by closing and site directed mutagenesis The vector backbone of the single-copy integration plasmid originates from the pNH605 plasmid was used for cloning ^45^. The pNH605 plasmid was digested with BamHI and NotI. For the WT plasmid construction, the digested pNH605 plasmid and the *UME6*^WT^ PCR fragment (∼700bp upstream of the START codon, and ∼700bp downstream of the STOP codon) were assembled. For the *UME6*^T99N^ the digested pNH605 plasmid and the overlapping PCR fragments with point mutation were assembled. For generating the *RIM11*^Y199F^ mutation, the genomic *RIM11* locus was first C-terminally tagged with mNeonGreen (mNG). The *RIM11*-mNG locus, including ∼700bp upstream of the START codon, was amplified and assembled with the digested pNH605 plasmid (*pNH605-RIM11*). The *pNH605-RIM11* was amplified by making use of the Q5® Site-Directed Mutagenesis Kit (New England Biolabs) following the instructions of the manufacturer to generate *pNH605-RIM11* ^Y199F^. The *IME1*^L321F^ plasmid was generated by site directed mutagenesis using pNH604-pIME1-sfGFP-IME1 described previously as a template. The plasmids were verified by sequencing. The list of the yeast strains and plasmids used in Table S2 and S3. *UME6* was C-terminally tagged with the TurboID tag including 3 tandem copies of the Myc epitope (TurboID-3xMyc)^46^.

### Growth conditions

For a standard meiosis time-course, cells were inoculated in liquid YPD (1% yeast extract, 2% peptone, 2% glucose, 22.4 mg/L uracil, and 80 mg/L tryptophan) grown to saturation overnight. The cells were transferred to liquid BYTA media (buffered, yeast extract, tryptone, acetate) OD_600_ =0.4. After 16-18h the cells were pelleted and washed once with sterile water before being transferred to liquid sporulation media (SPO) OD_600_ 2.5. For the mid-log phase samples, cells were inoculated in YPD, grown overnight and then diluted back to OD_600_ = 0.5. the next morning. The samples were taken when the cells reached OD_600_ =1.

To inhibit the TORC1, the cells were grown to mid-log phase OD_600_ 1, as described in the standard meiosis protocol. The cells were then treated with 1μM rapamycin (Sigma-Aldrich, R0395). To inhibit the PKA pathway, an ATP analogue-sensitive strain, *tpk1-as*, was used^8^. The ATP analogue 1NM-PP1 was used to inhibit PKA activity (Calbiochem, Merck Millipore). Cells were grown to mid-log phase OD_600_ 1, and subsequently cells were treated with 5μM of 1NM-PP1.

To induce expression from *CUP1* promoter, cells were grown to 16h BYTA. At this stage, the cells are treated with 50μM CuSO4 (ThermoFisher) for 2h. In SPO the cells were treated again with 25μM CuSO4.

For the Ume6-TID experiment, cells were grown to saturation overnight and diluted back to 50ml fresh YPD OD_600_ = 0.25. OD_600_ = 0.5 the cells were treated with 50μM biotin (Sigma-Aldrich, B4501) and harvested when the cultures reached OD_600_ =1. For sporulation conditions, cells were grown following the standard meiosis protocol. Cells were transferred to 50ml SPO OD_600_ =1.8 and treated immediately with 50μM biotin. At 4h SPO, cells were harvested.

For mitotic Sin3 depletion assays using auxin induced degron *SIN3-AID*, cells were grown in YPD. After overnight incubation, cultures were back diluted to an OD_600_ = 0.2 and grown for ∼3 h to exponential phase (OD_600_ ≥0.5). Once exponential phase was reached (0 hours), all cultures were treated with CuSO4 (50 µM final) and 3-indoleacetic acid (IAA; 200 µM final).

### Western blotting

For Western blot analysis, 5 OD_600_ units were pelleted and resuspended in 1 ml 5% TCA and put into the fridge for at least 10min. Cells were then pelleted and resuspended in 1ml 100% acetone and centrifuged at 13,200 rpm for 5min. The pellets were dried for 1.5h at RT before being resuspended in 100μl protein breakage buffer (50mM Tris pH 7.5, 1mM EDTA supplemented with 2.7mM DTT). 100μl of glass beads (Ø 0.5mm, BioSpec) was added and the cells were lysed in the bead-beater for 3min. 50μl of 3x SDS loading buffer (187.5 mM Tris pH 6.8, 5.7% β-mercaptoethanol, 30% glycerol, 9% SDS, and 0.05% Bromophenol Blue) was added to the protein lysates and boiled for 5min at 95°C.

The WB samples were loaded on a 4-20% or 7% Midi CriterionTM TGXTM Precast gel (BioRad) and ran at 180V for 45min in 1x running buffer (190mM glycine, 25mM and 3.4mM SDS). Trans-Blot SD Semi-Dry Transfer Cell (Bio-Rad) or the Trans-Blot Turbo Transfer System (Bio-Rad) were used for transfer to membranes. The former one was performed in a semi-dry transfer buffer (48mM Tris Base, 39mM glycine, 0.04% SDS and 10% methanol) onto a methanol-activated PVDF membrane. The latter one in 1x of commercially available transfer buffer (BioRad) onto a nitrocellulose membrane. The membrane blocked (1% BSA and 1% non-fat powdered milk in 1x phosphate-buffered saline (PBS) buffer supplemented with 0.01% Tween-20 (0.01% PBST), washed 3x in 0.01% PBST buffer and incubated with the primary antibodies added to the blocking buffer overnight at 4°C.

Hexokinase (Hxk1) was used as the loading control for WB in this thesis. The list of antibodies can be found in Table 3. The next day, the membrane was washed three times with 0.01% PBST buffer and incubated with secondary antibodies. If images were taken with the Li-COR Odyssey® CLx, secondary antibodies IRDye 800CW anti-mouse and IRDye 680RD anti-rabbit (LI-COR) were used. Anti-mouse and anti-rabbit IgG horseradish peroxidase (HRP)-linked antibodies (Amersham) and streptavidin HRP-conjugated antibodies (Sigma-Aldrich) were used for ECL detection Reagents (Sigma-Aldrich) and scanning on the Amersham Imager 600 Instrument (GE Healthcare). Image processing and quantification was done with the Image StudioTM Lite software.

### RT-qPCR

Total RNA was extracted from 5 OD_600_ units using TES buffer (10mM Tris HCl pH 7.5, 10mM EDTA (pH8), 0.5% SDS) and acid-phenol:chloroform (Ambion) at 65°C, followed by ethanol precipitation, DNAse treatment (rDNAsa), and column-purified using NucleoSpin RNA kit (Macherey-Nagel). 500ng of total RNA was used for reverse transcription using ProtoScript II First Strand cDNA Synthesis Kit (New England BioLabs). For quantification, the QuantStudioTM 7 Real-Time PCR. The gene expression values are relative to the housekeeping gene expression *ACT1*.

For Figure 6, 5 µg of purified total RNA was then treated with DNase (TURBO DNA-free kit, Thermo Fisher (MA, USA) according to manufacturer, and 4 µL (<1 µg) of DNase treated total RNA was then reverse transcribed into cDNA with the use of random hexamers (Superscript III Supermix, Thermo Fisher) according to manufacturer’s instructions. cDNA was then quantified using the SYBR green mix (Life Technologies (CA, USA)) and measured using the Applied Biosystem StepOnePlusTM Real-Time PCR system (Thermofisher – 4376600). Signald were normalized to *ACT1*. PCR primers used are listed in Table S4.

### Microscopy analysis

Images were taken with an Eclipse Ti-E inverted microscope system (Nikon) using the 100x oil objective. Proteins tagged with mNeonGreen (mNG) and superfolder GFP (sfGFP) were imaged with a GFP filter with an exposure time of 500msec and proteins tagged with GFP with 1sec. Proteins tagged with mCherry were imaged with a mCherry filter with an exposure time of 100ms. Image analysis was carried out using Fiji, where whole cell selection was determined manually and nuclei selection was determined via segmentation^47^. The mean gray values (mean intensity) were calculated by taking the sum of the gray values of all pixels within the selected object over the number of pixels. To investigate on nuclear localisation of proteins, the ratio between the nuclear mean intensity over the whole mean intensity was taken.

### DAPI

Cells were resuspended in 1μg/mL 4’,6-diamidino-2-phenylindole (DAPI) in 1x phosphate-buffered saline (PBS) buffer and visualised by fluorescent microscopy. 200 cells were counted per sample where each cell was assessed for the number of nuclei. Cells with one DAPI mass were considered no meiosis. Cells with two or more DAPI masses were considered meiosis. The percentage of cells that underwent the first and second meiotic division over the total number of cells was calculated.

### TurboID MS

Cells were pelleted, washed two times in sterile water, and four washes in RIPA buffer 0.4% SDS (50mM Tris HCL pH 7.5, 150mM NaCl, 1.5mM MglCl2, 1mM EGTA, 0.04%SDS, 1% NP-40). Cells were resuspended in 2ml 5% TCA per 5 OD_600_ units incubated 4°C overnight, pelleted, washed in 100% acetone, resuspended in in TE buffer (100mM Tris pH 7.5, 1mM EDTA) supplemented with 2.7mM DTT, per 18OD units. 500μl of zirconia/silicate beads (Ø 0.5mm) were added and the cells were lysed in the bead-beater. 3xSDS sample buffer (187.5mM Tris pH 6.8, 30% glycerol, 3% SDS) was added and the samples were boiled. For affinity purification, 400μl of streptavidin M280 beads (ThermoFisher) were used per sample, incubated at 4°C for 3h. Beads were washed once with wash buffer, four times with RIPA 0.4% SDS buffer and five times with 20mM ammonium bicarbonate. Beads were re-suspended in 50μl of 50mM ammonium bicarbonate.

Samples were prepared using on-beads tryptic digestion protocol prior to mass spectrometry. Samples were reduced with 5mM DTT for 1 h at 37 °C in a thermomixer. Proteins were then alkylated with 10 mM IAA for 30 min in the dark at room temperature. After reduction/alkylation Beads were trypsinised in order to digest biotinylated proteins (0.4ug/ul trypsin in 50mM NH4HCO3). Digestion was performed overnight at 37 °C in a thermomixer at 450 rpm. The reaction was stopped with 10% formic acid (FA). Peptides were recovered and the beads were discarded utilising a magnetic rack. Finally, a C18 clean-up was performed using EV2018 EVOTIP PURE. The desalting was done according to manufacturer’s instructions protocol. Briefly, the Evotips tips were conditioned with 0.1% Formic acid in acetonitrile; equilibrated with 0.1% Formic acid in water and approximately 1 µg of each digested sample was loaded onto Evotips. Peptides were eluted from the Evotips with 50% acetonitrile into a vial and vacuum dried by SpeedVac to remove any traces of organic solvents. Finally, the dried peptides were re-suspended in 0.1% formic acid.

### Mass spectrometry analysis

The resulting peptides were analysed by nano-scale capillary LC-MS/MS using an Ultimate U3000 HPLC (ThermoScientific Dionex, San Jose, USA) to deliver a flow of approximately 300 nL/min. A C18 Acclaim PepMap100 5 µm, 75 µm x 20 mm nanoViper (ThermoScientific Dionex, San Jose, USA), trapped the peptides prior to separation on an EASY-Spray PepMap RSLC 2 µm, 100 Å, 75 µm x 500 mm nanoViper column (ThermoScientific Dionex, San Jose, USA). Peptides were eluted with a 90 min gradient of acetonitrile (2%v/v to 80%v/v). The analytical column outlet was directly interfaced via a nano-flow electrospray ionisation source, to a hybrid quadrupole orbitrap mass spectrometer (Lumos Tribrid Orbitrap mass spectrometer, ThermoScientific, San Jose, USA). Data dependent analysis was carried out, using a resolution of 120,000 for the full MS spectrum, followed by MS/MS spectra acquisition in the linear ion trap using “TopS” mode. MS spectra were collected over a m/z range of 300–1800. MS/MS scans were collected using a threshold energy of 32% for collision induced dissociation.

### Turbo-ID data analysis

LC-MS/MS raw files were processed in MaxQuant (version 2.0.3.1). The LFQ algorithm and match between runs settings were selected. Data were searched against the reviewed UniProt Saccharomyces cerevisiae proteome using the Andromeda search engine embedded in MaxQuant.

Trypsin was set as the digestion enzyme (cleavage at the C-terminal side of lysine and arginine amino acid residues unless proline is present on the carboxyl side of the cleavage site) and a maximum of two missed cleavages were allowed. Cysteine carbamidomethylation was set as a fixed modification, while oxidation of methionine and acetylation of protein N-termini were set as variable modifications. The “match between runs” feature was used with a matching time window of 0.7 min and an alignment time window of 20 min. Label-free quantification was performed using the MaxLFQ feature included in MaxQuant according to default LFQ parameters. Minimum peptide length was set at 7 amino acid units. FDR, determined by searching a reverse sequence database, of 0.01 was used at both the protein and peptide level.

The MaxQuant protein groups output file was imported into Perseus software (version 1.4.0.2) for further statistical analysis and data visualization. Contaminant and reverse protein hits were removed. LFQ intensities were log2 transformed. Missing values (NaN) were imputed from a normal distribution with default values. For each sample, the triplicates were grouped. Data were filtered for at least two out of three replicates LFQ intensity values in at least one group. Protein LFQ intensities were normalised and missing values were imputed by values simulating noise around the detection limit using the default parameters. A protein was considered significantly differentially expressed when FDR < 0.05.

### Microfluidic cell culture

Y04C CellASIC microfluidic devices (http://www.cellasic.com/) were used for cell culture at 25°C, with 0.6 psi isobaric flow rate. To promote meiosis entry, cells were grown in yeast peptone dextrose (YPD) at 30°C in an orbital shaker at 295 rpm for 24 hours; once OD_600_ reached ∼2-3, 50 µl of cells were sonicated for 4-6 seconds at 3 W, and loaded in the microfluidic device using 1-2 pulses at 20 psi for 5 seconds. Meiotic induction was triggered by exposing the cells for at least 20 hours to sporulation medium (SPO) composed of 0.6% Potassium Acetate, 2 % sorbitol, 40 mg/L Adenine, 40 mg/L Uracil, 20 mg/L Histidine, 20 mg/L Leucine, 20 mg/L Tryptophan, 0.02 % Raffinose (from a 20 % w/v stock solution) and pH adjusted to 8.5 using a 0.25 M sodium carbonate solution. For return to growth experiments, cells were transferred to synthetic complete medium (SCD: 1% succinic acid, 0.6% sodium hydroxide, 0.5% ammonium sulfate, 0.17% yeast nitrogen base without amino acids or ammonium sulfate, 0.13% amino acid dropout powder (complete), 2% glucose) after 24 hours of microfluidic exposure to SPO at 25°C maintaining a flow rate of 0.6 psi.

### Time-lapse microscopy

Microfluidics experiments were performed using at least two different microscopy set ups for reproducibility. Both set ups used an automated Zeiss Observer Z1 microscope controlled by ZEN pro software (Zeiss), images were acquired at 12 min sampling rate using an 40X Zeiss EC Plan-Neofluar 40X 1.3 NA oil Ph 3 M27 immersion objective. Image focus was controlled using Definite Focus 2.0 or 3.0. Images were registered using an AxioCam HR Rev 3 camera or an AxioCam 712 monochrome. At least 5 fields of view were imaged for each biological replicate. Light sources used were COLIBRI 2.0 light-emitting diodes (LED) or an X-CITE XYLIS XT720S lamp (Excelitas Technologies). Sequential non-phototoxic imaging of four minimally cross-talking fluorophores during meiotic development was achieved through tailored dichroic mirrors and bandpass filters using the following fluorophores: Teal-cyan fluorescent protein (mTFP1), green-yellow fluorescent protein (mNeonGreen, Superfolder GFP (sfGFP), mKusabira-Orange κappa fluorescent protein (mKOκ), eqFP611 red fluorescent protein variant (mRuby3), mRFP1.5-derived red fluorescent protein (mCherry) and phase-contrast^48–53^. Spectral unmixing of strains containing red and orange fluorophores (FK10696 and FK11243) created images were the final mKOκ image was equal to the original mKOκ image minus 0.51 the red image.

### Image processing and quantification of cellular features

Image analysis code is available at https://github.com/alejandrolvido/Quiescence-Entry and https://github.com/alejandrolvido/Spectral-Imaging. Images were acquired using a ZEN pro software (Zeiss) with a 2 × 2 binning configuration, in uncompressed TIFF format. Images were converted to double format and de-noised using the functions double(), medfilt2(). Background-correction and cell segmentation was done according to ^54^. A Gaussian fit method was used to define the nuclear area of single cells according to (Arguello-Miranda et al., 2018). Nuclear and cytoplasmic concentrations were obtained using the function Get_Sphere_Vol(), which labels the objects within a cell mask, computes the equivalent diameter for each object based on its pixel area, and calculates the total nuclear or cytoplasmic volume.

### Time series and clustering analyses

The presence of the first meiotic nuclear division, which marks the point of no-return in the meiotic program, was used to classify cells as meiosis or no meiosis ^55^. To obtain MI times, the first derivative of the fluorescence intensity times series of a nuclear marker, such a histone Htb1-mCherry or IME1pr-mRuby3, was calculated and the point with the highest inflection was assessed as the first meiotic division, this assignment was manually confirmed using custom MATLAB code to score the onset of MI nuclear division by directly inspecting fluorescent image time series ^54^. Rim11 peaks were determined as the highest intensity point in the Rim11-mNG or Rim11-mKOκ nuclear fluorescence intensity time series over a window of 60 time points before the time of MI nuclear division. For locating Ume6-mTFP1 and sfGFP-Ime1 peaks, the time window was restricted to 40 time points before the first meiotic nuclear division.

Single cell meiotic time series clusters were created by applying the function kmeans() to the first 80 time points after starvation onset using correlation distance, 10 replicates, a maximum iteration limit of 10000, and with an experimentally defined number of clusters.

### Statistical analyses

Statistical analyses were performed using GraphPad Prism Version 9.0.0 for Windows, GraphPad Software, www.graphpad.com. One-way or two-way ANOVA tests were carried out, depending on the number of conditions tested.

Statistical analyses were conducted in MATLAB, treating each independent microfluidic device as a biological replicate. Kolmogorov-Smirnov tests (K-S test) determined statistical significance set at p < 0.05 using the function kstest2(). Line plots correspond to the biological replicates average surrounded by the 95 % confidence intervals, represented as shaded areas. Scatter plots depicted R-squared values calculated using corrcoef().

## Acknowledgement

We thank the Frank Uhlmann and Katrin Rittinger for critical reading of the manuscript. This research was funded in whole, or in part, by the Wellcome Trust (FC001203). For the purpose of Open Access, the author has applied a CC BY public copyright licence to any Author Accepted Manuscript version arising from this submission. This work was supported by the Francis Crick Institute (FC001203), which receives its core funding from Cancer Research UK (FC001203), the UK Medical Research Council (FC001203), and the Wellcome Trust (FC001203). ACSJ is supported by the Eric and Wendy Schmidt AI in Science Postdoctoral Fellowship, a Schmidt Futures program.

---

**Figure S1.**
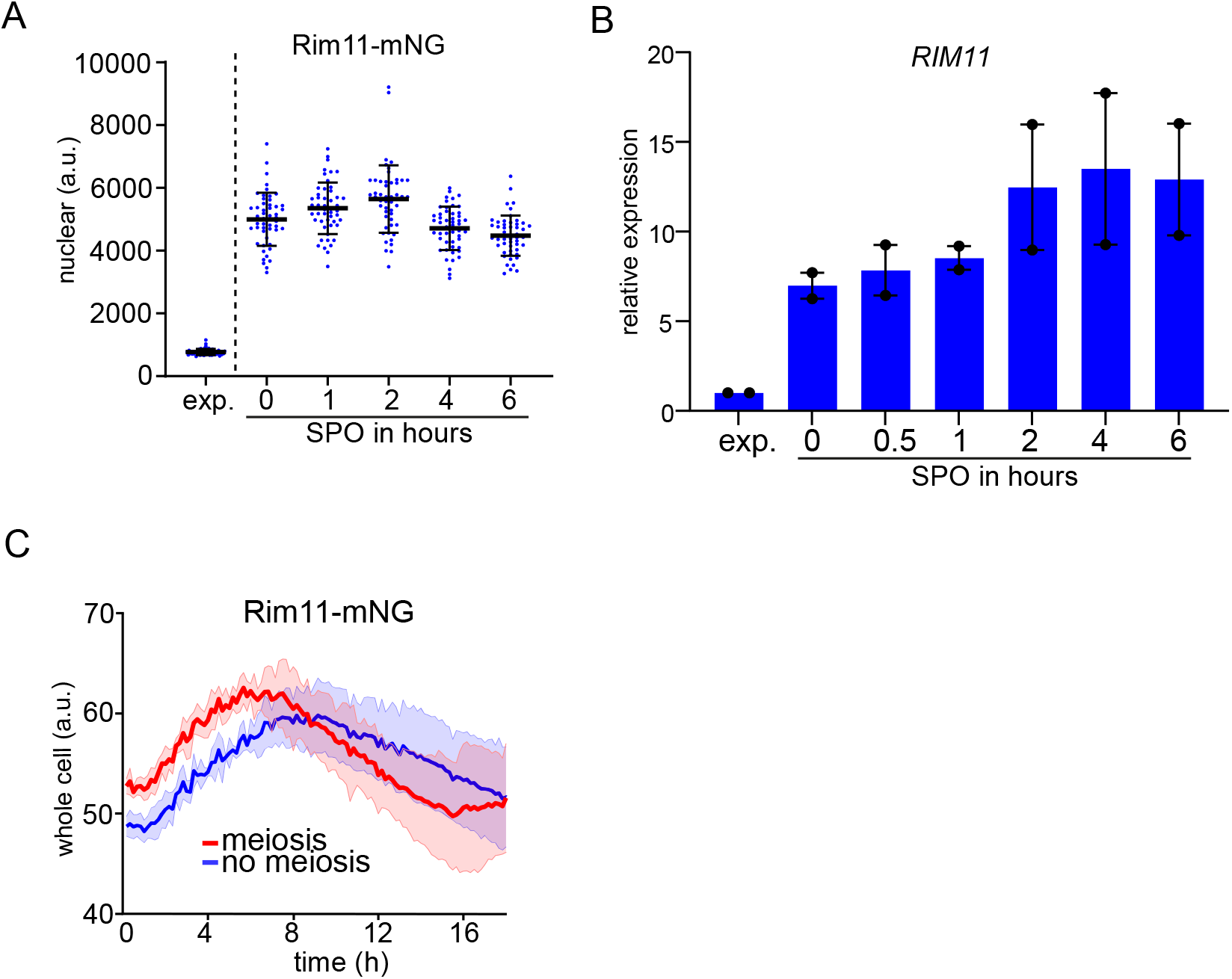
Rim11 expression and localization dynamics prior and during meiosis. **(A)** Nuclear concentrations of Rim11 in exponential growth (exp.) and in cells entering meiosis. We used a strain harbouring epitope tagged Rim11 with mNeongreen (Rim11-mNG) and histone H2B epitope tagged with mCherry (Htb1-mCh) (FW10297). **(B)** *RIM11* expression determined by RT-qPCR in wild-type cells (FW1511) grown untill exponential phase (exp.) and cells induced to enter meiosis. Samples were normalized to the time point 0 hours. n=2 biological repeats were performed. The *RIM11* signals were normalized over *ACT1*. **(C)** Quantification of whole cell concentrations of Rim11 using live cell imaging set up using the strain described in A. Indicated are the mean traces of cells that underwent meiosis (meiosis) and cells that did not undergo meiosis (no meiosis). Shown are the Rim11 cytoplasmic concentrations.

**Figure S2.**
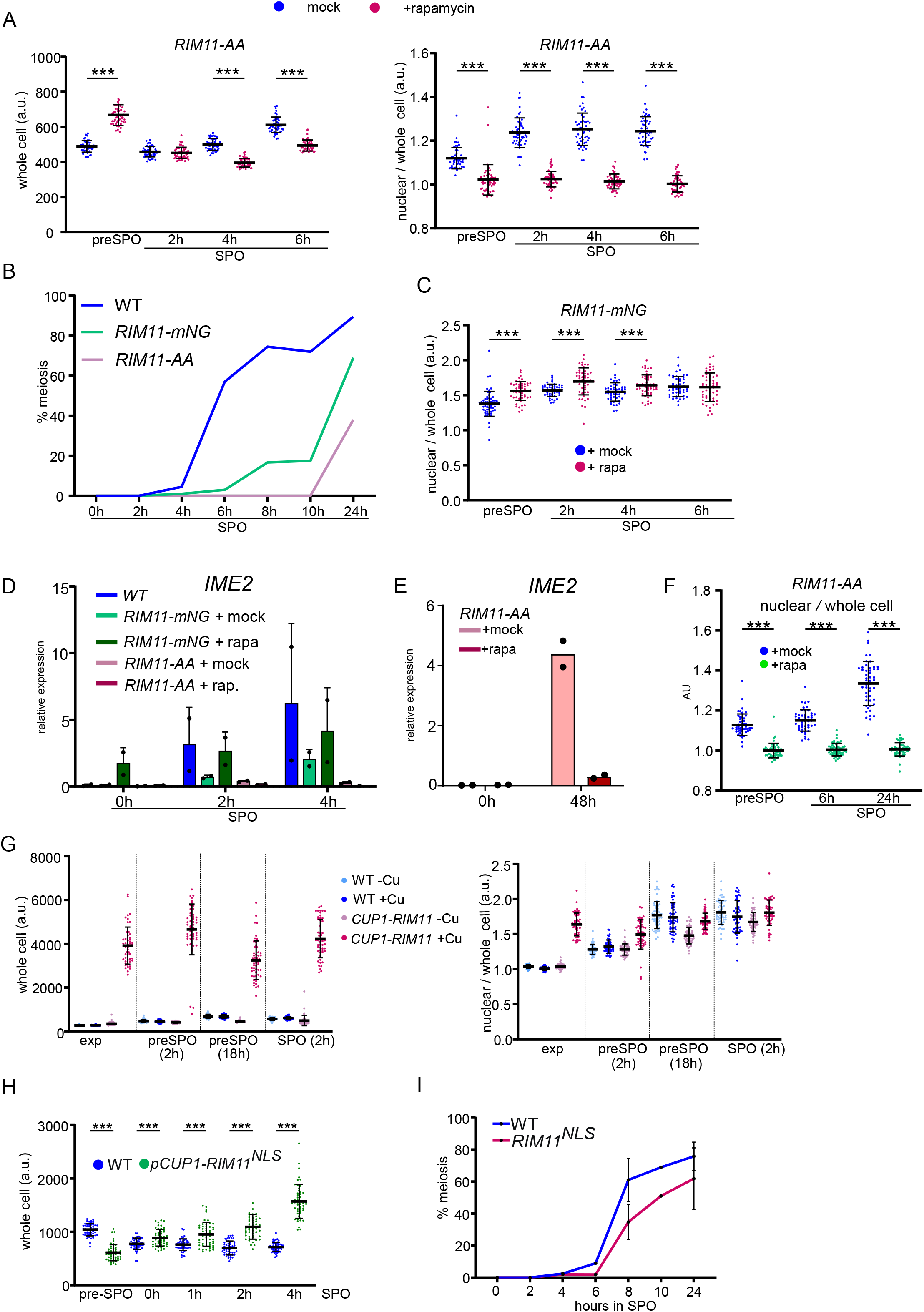
Rim11 nuclear localization is required for meiosis, but not sufficient. **(A)** Quantification of whole cell signal (left) and nuclear over whole cell signal (right) of Rim11-AA in either mock treated or treated with rapamycin (FW11124) in pre-sporulation medium (pre-SPO), and different time points in sporulation medium (SPO). **(B)** Onset of meiosis in WT (1511), Rim11-nNG (FW11126), Rim11-AA (FW11124). Cells were induced to sporulate, and samples were taken at the indicated time points, fixed, and stained with DAPI. Cells that contained two more DAPI masses were considered to have undergone meiosis. **(C)** Similar analysis as A, except that nuclear over whole cell signals for the Rim11-mNG (FW11126) was determined. **(D)** *IME2* expression in strains described in A-C in cells induced in enter meiosis. Samples were taken at the indicated time points, RT-qPCR was performed to determine *IME2* expression. The relative signals normalized over *ACT1* of n=2 biological repeats are shown. **(E)** Similar as D, except that *IME2* expression was determined in Rim11-AA (FW11124) after 24 hours in SPO in mock and rapamycin treated cells. **(F)** Nuclear over whole cell signal of Rim11-AA in mock and rapamycin treated cells induced to enter meiosis. **(G)** Whole cell signal (left) and nuclear over whole cell signal (right) of Rim11 when expressed from *CUP1* promoter in cells induced to enter meiosis (FW10297, FW11012). **(H)** Whole cell signal of WT Rim11 (FW10776) and Rim11 fused to NLS under control of *CUP1* promoter (FW11206) in cells induced to enter meiosis. **(I)** Onset of meiosis in WT (FW1511) and *pCUP1-NLS-RIM11* (FW11206). Cells were induced to sporulate, and samples were taken at the indicated time points, fixed, and stained with DAPI. Cells that contained two more DAPI masses were considered to have undergone meiosis.

**Figure S3.**
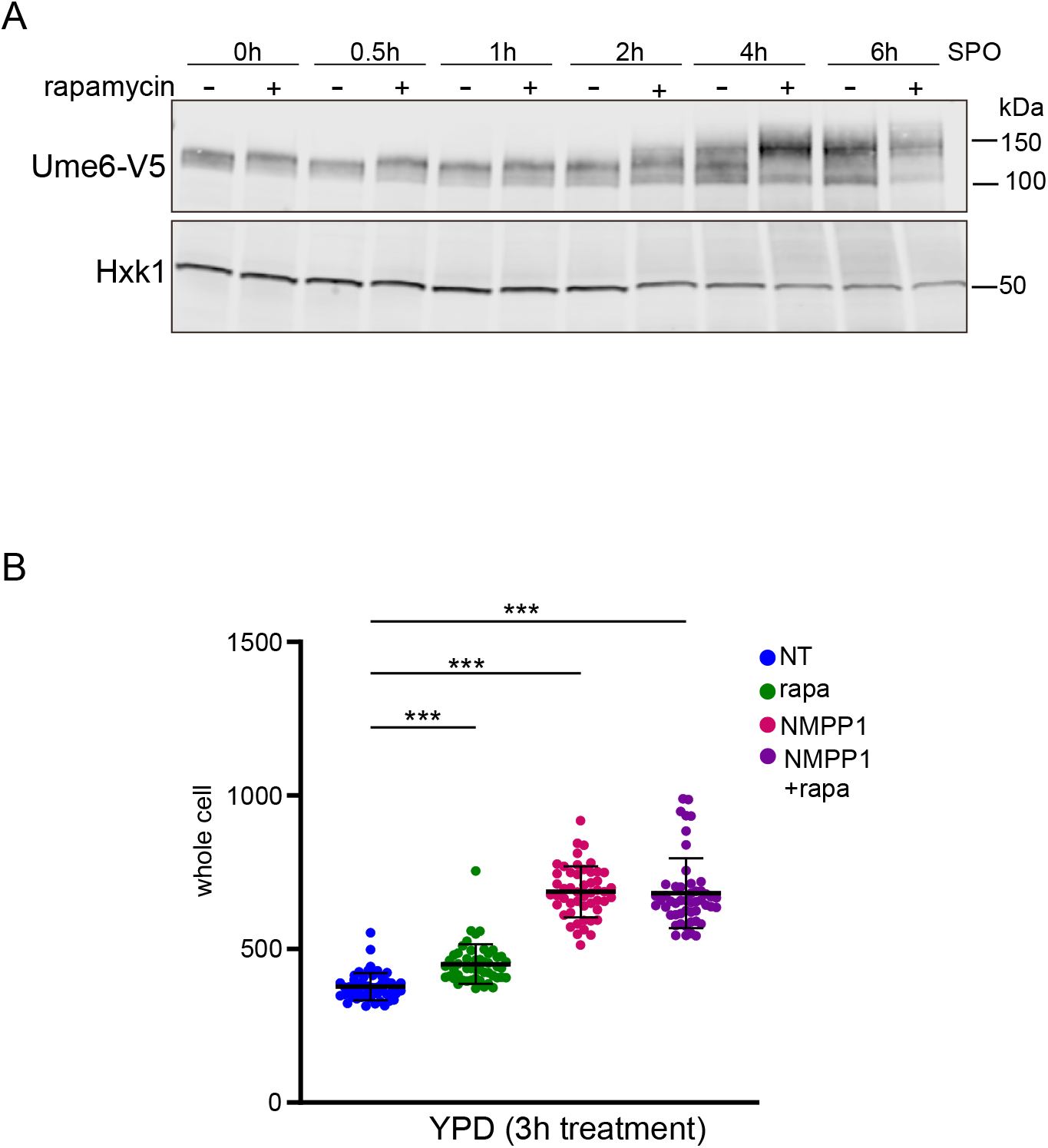
TORC1 and PKA control Rim11 localization and expression. **(A)** Ume6 migration as determined by western blotting (FW1208). Cells were induced to enter meiosis and were either untreated or treated with rapamycin. Western blot membrane was also probed anti-Hxk1 antibodies as a loading control Hxk1. **(B)** Rim11-mNG whole cell signal in the *tpk1-as* allele background (FW10483). Cells were grown in rich medium and were untreated (NT), or treated with rapamycin, NMPP1 or both compounds.

**Figure S4.**
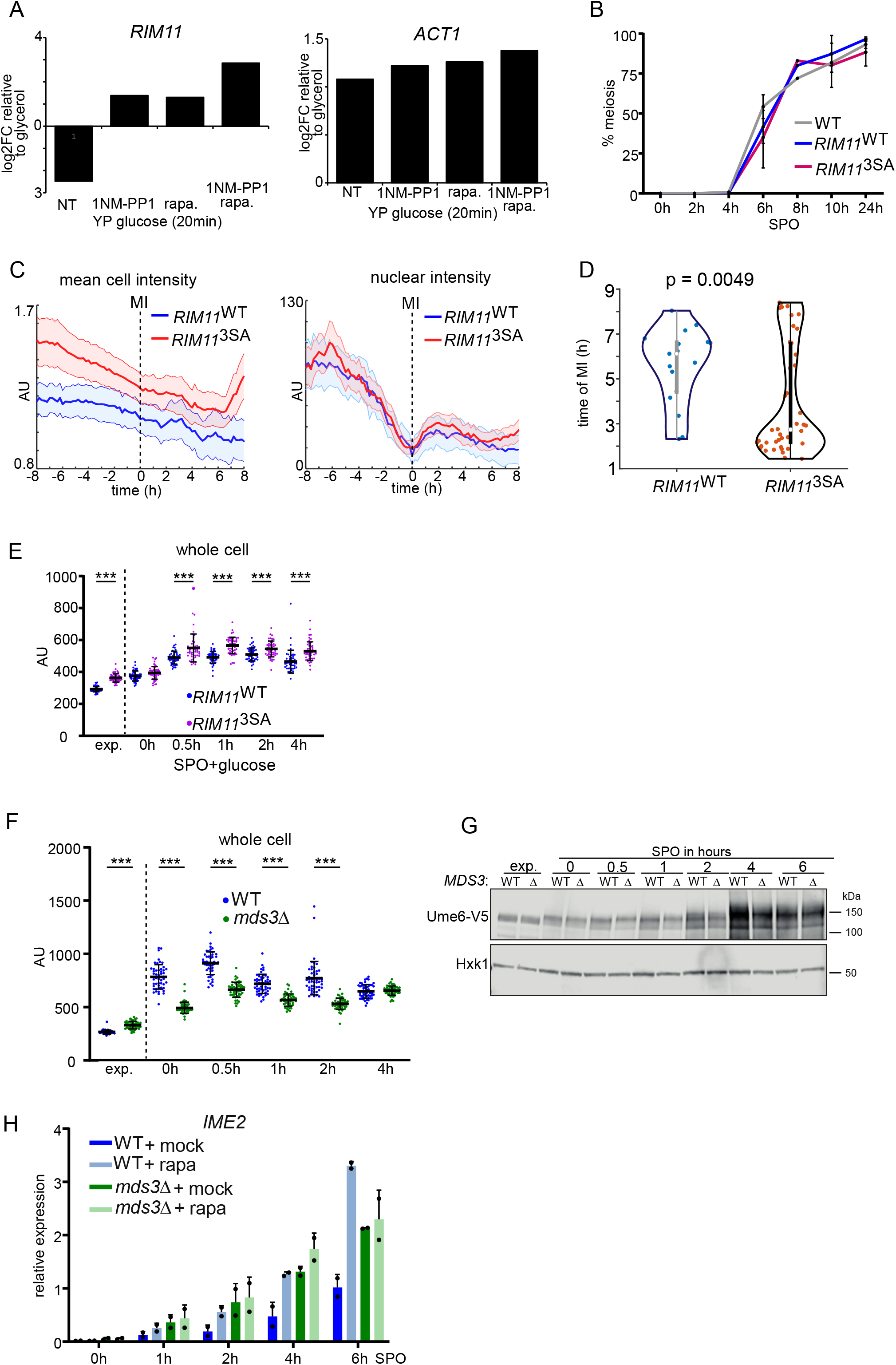
PKA and TORC1 control Rim11 via distinct mechanisms. **(A)** *RIM11* and *ACT1* expression comparing YP glycerol to a shift to YP glucose in the PKA^AS^ allele background. Cells were either not treated (NT), or treated with 1NM-PP1, rapamycin or both compounds for 20 minutes. Shown are log2 fold change (FC) for *RIM11* and *ACT1*. The data were taken from ^25^. **(B)** Onset of meiosis of Rim11^WT^ and Rim11^3SA^ (FW1511, FW10776 and FW10778). The mean signals of n=3 repeats are shown. **(C)** Quantification of live cell imaging of Rim11^WT^ and Rim11^3SA^. Shown are whole cell concentration (left) and nuclear intensity (right). Time points were aligned according to the MI division. **(D)** Same experiment as in B but showing the timing of meiosis in Rim11^WT^ and Rim11^3SA^. **(E)** Whole cell quantification of Rim11^WT^ and Rim11^3SA^ in SPO + 1% glucose. **(F)** Whole cell quantification Rim11-mNG in WT and *mds3*Δ in cells induced to enter meiosis. **(G)** Ume6 migration as determined by western blotting in WT, *mds3*Δ, and *rim11*Δ. Cells were induced to enter meiosis and samples were taken at the indicated time points. Membranes were probed with anti-V5 antibodies and Hxk1 antibodies as a loading control (FW1208, FW11251). **(H)** *IME2* expression in cells induced to enter meiosis in WT and *mds3*Δ that were either untreated or treated with rapamycin (FW10297 and FW10718). *IME2* expression signals were normalized over *ACT1*. The mean signals of n=2 biological repeats are shown.

**Figure S5.**
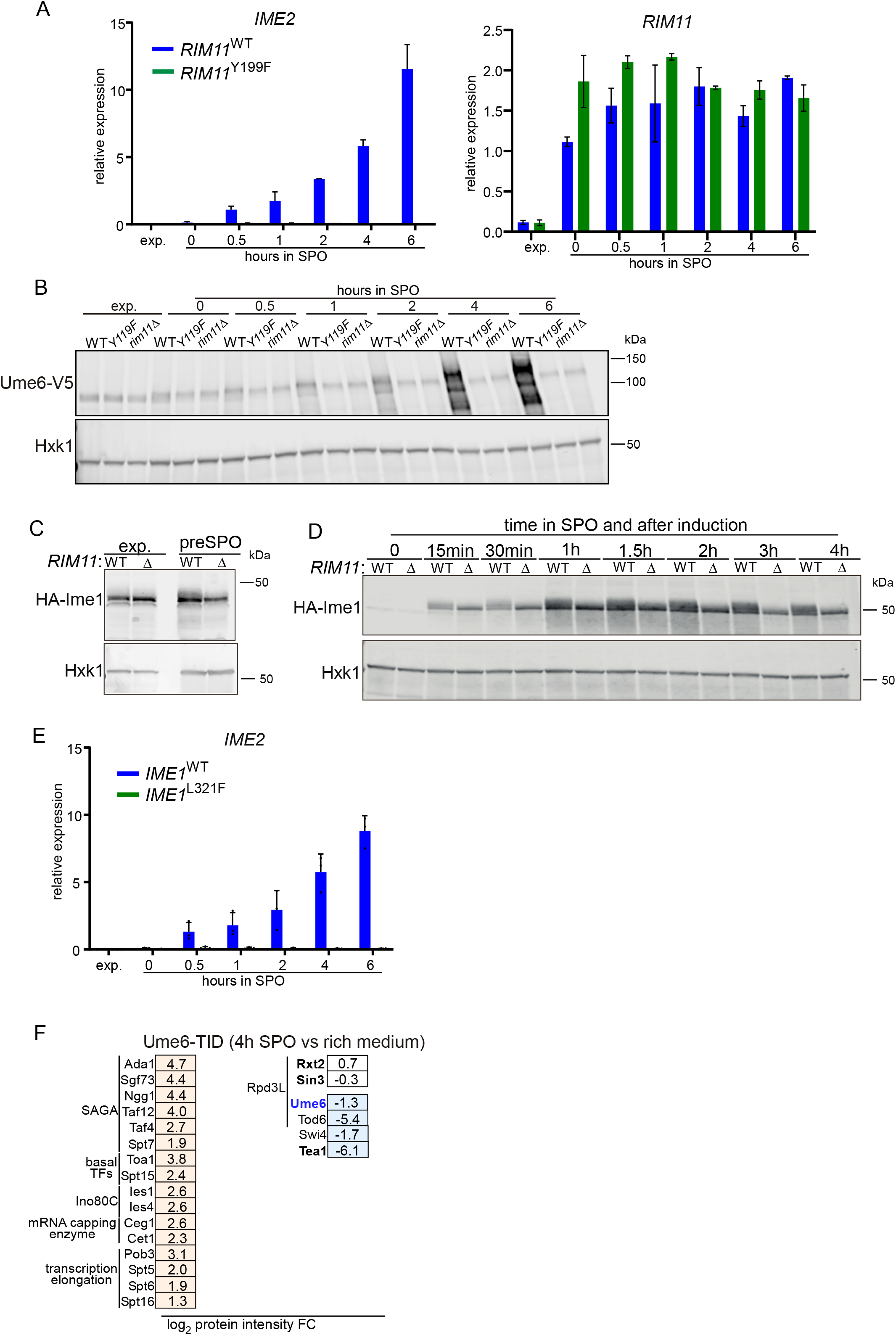
Ime1 is essential for Rim11 directed Ume6 phosphorylation. **(A)** *IME2* and *RIM11* mRNA expression in Rim11^WT^ and Rim11^Y199F^ (FW10776, FW10983) cells induced to enter meiosis. mRNA expression levels were normalized to *ACT1*. The mean value of n=3 biological repeats is shown. **(B)** Ume6 expression and migration as determined by western blotting in *Rim11^Y119F^* and *rim11*Δ cells induced to enter meiosis (FW1208, FW11097, FW10033). Membranes were probes with anti-V5 antibodies and Hxk1 antibodies. **(C-D)** Ime1 expression and migration in exponential growth, and in cells induced to enter meiosis. Ime1 tagged with HA was expressed from the *CUP1* promoter in induced in WT and *rim11*Δ cells (FW2444, FW10373). **(E)** *IME2* expression in *IME1*^WT^ and *IME1*^L321F^ (FW11231, FW11233). *IME2* expression was normalized to *ACT1*. The mean value of n=3 biological repeats is shown. **(F)** TurboID analysis of Ume6 (Ume6-TID) comparing 4h SPO to rich medium conditions (FW11422). Log2 protein intensity fold change (FC) comparing 4h SPO to rich medium. Proteins involved in transcription or known to interact with Ume6 are shown.

**Figure S6.**
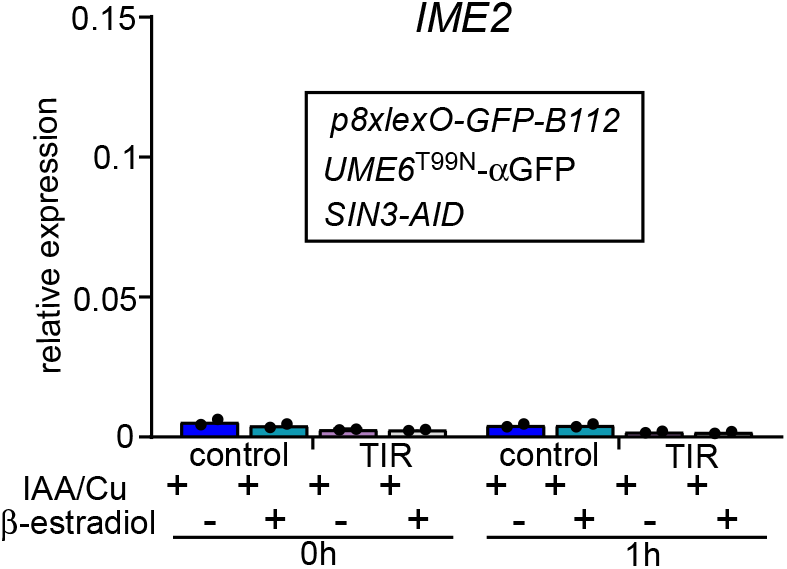
Rewiring of Ume6 regulon makes Rim11 and Ime1 dispensable. *IME2* mRNA expression in cells grown in exponential growth phase. Cells expressing Ume6^T99N^-aGFP and GFP-B112 under control of eight lexO sites (*p8xlexO-GFP-B112*) and Sin3-AID were untreated or treated with β-estradiol (UB36868, UB36874). Additionally, all cells were treated with 3-indoleacetic acid (IAA) and copper sulphate. The mean signal of n=2 biological repeats is shown.

**Figure S7.**
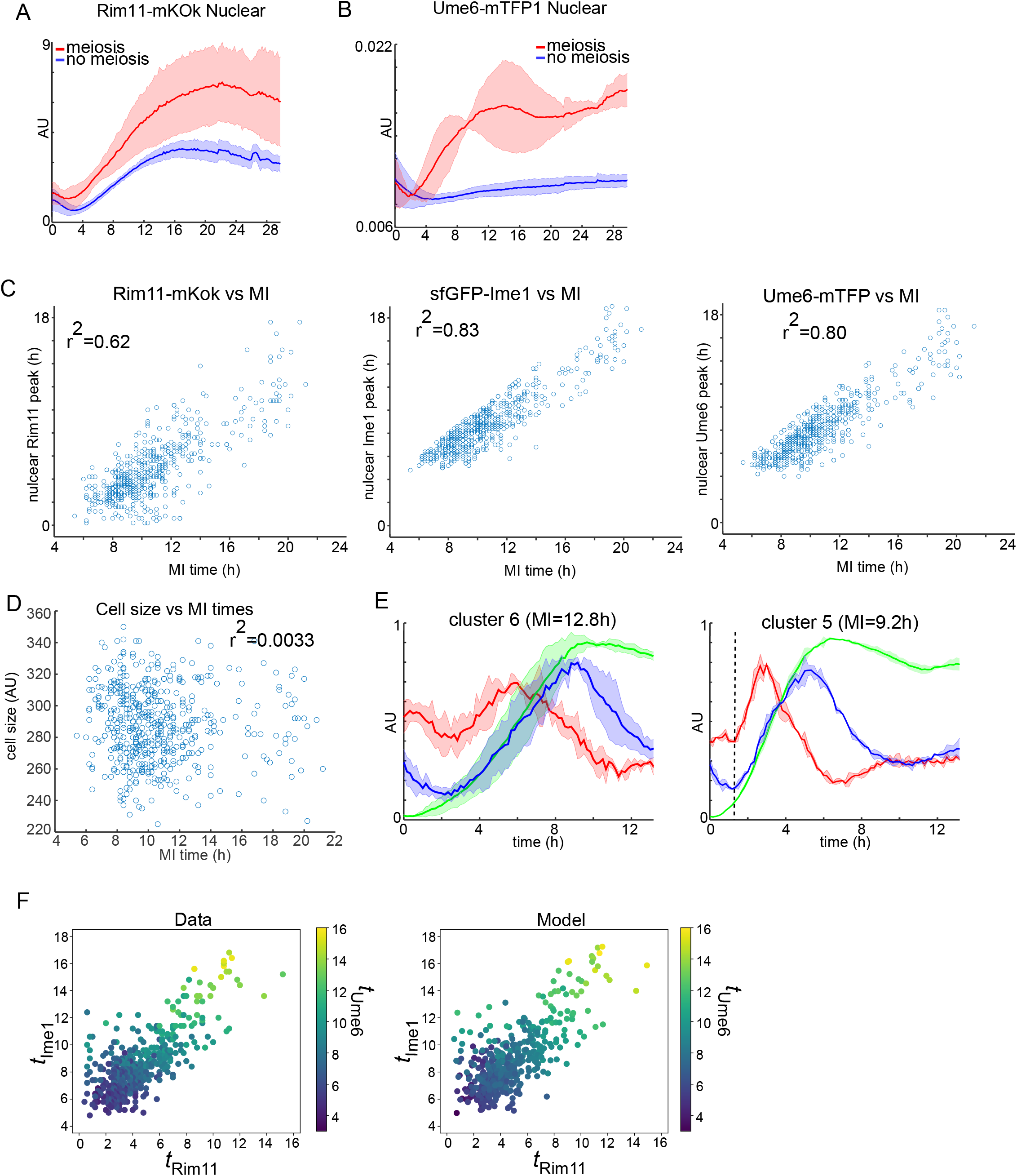
Single cell analysis and modelling reveals the timing dynamics Rim11, Ime1, Ume6 and meiosis. Live cell imaging of the dynamics of Rim11, Ime1, Ume6 in meiotic and non-meiotic cells. We generated a strain with sfGFP-Ime1, Ume6-mTFP, and Rim11-mKok, and pIME1-NLS-mRuby3 (FW11243). **(A)** Nuclear concentrations of Rim1-mKok in meiotic and non-meiotic cells. **(B)** Nuclear concentrations of Rim11-mTFP in meiotic and non-meiotic cells. **(C)** Scatter plot showing the single cell data comparing timing of Rim11-mKok peak versus MI (left), sfGFP-Ime1 peak versus MI (middle), Ume6-mTFP peak versus MI (right). **(D)** Scatter plot showing the single cell data comparing timing of cell size versus MI. **(E)** Mean traces of Rim11, Ime1, Ume6 nuclear intensity for cluster 6 (left), and 5 (right) described in Figure 7D. **(F)** Time of Ume6 peak as a function of Ime1 and Rim11 peak timings for the cells that enter meiosis. The left panel shows the results from the time series data, while the right panel illustrates our model predictions. Besides the peak time in Ime1 and Rim11, the model reproduces the amplitudes of the peaks in Ime1 and Rim11 as well as the final value of Rim11 in the single cell data. The plot contains 461 out of 524 cells – of the remaining 63 cells, most were excluded because the maximal values in Ime1 and Rim11 of the model lie outside the first 18 hours after t=0.

