## Supplementary file 3 for "Multi-signal regulation of the GSK-3β homolog Rim11 governs meiosis entry in yeast"

### Mathematical Model

We employ a system of ordinary differential equations (ODEs) to model the regulatory network that controls the entry into meiosis. Within this framework, we capture the impact of TORC1 and PKA into three starvation signals, called $S_1$, $S_2$ and $S_3$, that lead to the localisation of Rim11 and regulate the Ime1 and Rim11 levels. All three starvation signals are assumed to follow a sigmoidal behaviour and thus vary between 0 to 1. To get this we use a simple logistic growth model with delay:

$$\frac{dS_1}{dt} = \theta_1(1 - S_1)S_1, \text{ if } t \geq t_1, \text{ else } \frac{dS_1}{dt} = 0 \text{ and } S_1 = 0.01$$

where $\theta_1$ denotes a growth rate, and $t_1$ denotes any delay in the onset of starvation. Similar equations govern $S_2$ and $S_3$.

Our model strives to recover the main trends and properties of our single cell time series data. For simplicity, we assume that all Rim11 present in the nucleus ($\text{Rim11}_N$) plays an active role for phosphorylation, while the Rim11 in the cytosol ($\text{Rim11}_C$) is assumed to be an inactive reservoir. Moreover, we only consider and aim to recover the concentrations of Ime1 and Ume6 in the nucleus. Based on these assumptions, the concentration of the unphosphorylated Ime1 in the nucleus as a function of time ($t$) is governed by the following ODE:

$$\frac{d\text{Ime1}}{dt} = s_1 \cdot S_1 - d_1 \cdot \text{Ime1} - p_1 \frac{\text{Ime1} \cdot \text{Rim11}_\text{N}}{c_1 + \text{Ime1}} + u_1 \cdot \text{Ime1}_\text{P} + b_1.$$

Here, $s_1$, $d_1$, $p_1$, $c_1$, $u_1$ and $b_1$ denote model parameters: $s_1$ is connected to starvation, $d_1$ is a decay rate, $p_1$, $c_1$ are associated with phosphorylation and $u_1$ is associated to dephosphorylation of Ime1 and $b_1$ is a base transcription rate. The amount of unphosphorylated Ime1 is thus dictated by $S_1$ as well and its phosphorylated counterpart ($\text{Ime1}_P$) and thus by $\text{Rim11}_N$. Meanwhile, $\text{Ime1}_P$ can combine with Ume6 to form a complex (C). As we observe some adaptation to the stress in the level of Ime1, we introduce a decay term for $\text{Ime1}_P$ that depends on $S_1$ at an earlier time (that is at $t\text{-}k_1\tau_1$), this is a simple way of modelling an incoherent feedforward that is known to produce adaptive response:

$$\frac{d\text{Ime1}_\text{P}}{dt} = p_1 \frac{\text{Ime1} \cdot \text{Rim11}_\text{N}}{c_1 + \text{Ime1}} - u_1 \cdot \text{Ime1}_\text{P} - a_1 \cdot \text{Ime1}_\text{P} \cdot \text{Ume6} + a_2 \cdot C - d_2 \cdot \text{Ime1}_\text{P} \cdot S_1(t - k_1\tau_1)$$

Here, $\tau_1$ denotes the time at which Ime1 peaks for the mean single cell time series, while $a_1$, $a_2$, $d_1$, and $k_1$ are model parameters: $a_1$ and $a_2$ denote rates that describe the formation of the complex, and $d_1$ is a decay rate.

For Rim11, $S_2$ controls the concentration in the cytosol (modelling transcriptional regulation), while $S_3$ governs the localisation to the nucleus:

$$\frac{d\text{Rim11}_\text{C}}{dt} = s_2 \cdot S_2 - l_\text{cn} \cdot S_3 \cdot \text{Rim11}_\text{C} + l_\text{nc} \cdot \text{Rim11}_\text{N} - d_3 \cdot \text{Rim11}_\text{C} + b_2,$$

$$\frac{d\text{Rim11}_\text{N}}{dt} = l_\text{cn} \cdot S_3 \cdot \text{Rim11}_\text{C} - l_\text{nc} \cdot \text{Rim11}_\text{N} + l_\text{b} \cdot \text{Rim11}_\text{C} - d_4 \cdot \text{Rim11}_\text{N} \cdot S_3(t - k_2\tau_2).$$

Here, $l_{cn}$, $l_{nc}$ and $l_b$ are model parameters that are associated with localisation. Both, d3 and d4 are decay rates, while b2 is a basal transcription rate. Again, we have introduced a decay term that depends on the starvation signal ($S_3$) at an earlier time, at $t\text{-}k_2\tau_2$, based on the time ($\tau_2$) at which $\text{Rim11}_N$ peaks in the mean single cell time series, which is to produce an adaptive response to stress as observed in the data.

We have find Ime1 acts as an scaffold for Ume6 phosphorylation, so we assume complex C can get phosphorylated by $\text{Rim11}_N$ on Ume6 producing a new complex ($C_P$) of phosphorylated Ime1 and phosphorylated Ume6. The concentration of the complex C formed by $\text{Ime1}_P$ and Ume6 is given by

$$\frac{dC}{dt} = a_1 \cdot \mathrm{Ime1_P} \cdot \mathrm{Ume6} - a_2 \cdot \mathrm{C} - p_2 \cdot \frac{\mathrm{Rim11_N} \cdot \mathrm{C}}{c_2 + \mathrm{C}} + u_2 * \mathrm{C_P}$$

where $p_2$ and $c_2$ are model parameters that are associated with the phosphorylation and $u_2$ is a model parameter associated with dephosphorylation of the Ume6 in the complex. The concentration of this new complex is given by

$$\frac{d\mathrm{C_P}}{dt} = p_2 \cdot \frac{\mathrm{Rim11_N} \cdot \mathrm{C}}{c_2 + \mathrm{C}} - u_2 \cdot \mathrm{C_P}.$$

Finally, it is known that $\mathrm{C_P}$ regulates transcription of Ume6 promotor (in addition to the other early miotic genes, such as Ime2). So, the concentration of Ume6 is dictated by the formation of the complex C and the concentration of the phosphorylated complex through a positive feedback loop (based on the model parameters $p_3$, $c_3$ and $q$):

$$\frac{d\mathrm{Ume6}}{dt} = p_3 \cdot \frac{\mathrm{C_P^q}}{c_3^q + \mathrm{C_P^q}} - d_5 \cdot \mathrm{Ume6} + b_3 - a_1 \cdot \mathrm{Ime1_P} \cdot \mathrm{Ume6} + a_2 \cdot C,$$

Where $b_3$ is a base transcription rate and $d_5$ is a decay rate.

To recover the mean single cell time series qualitatively we tunes the parameters using a combination manual adjustment and fitting. We set $k_1$ to 0.85, $k_2$ to 0.55, $d_1$ to 0.22 per hour, $d_2$ to 3.0 per hour, $d_3$ to 18 per hour, $d_4$ to 0.53 per hour, $d_5$ to 3.0 per hour, $p_1$ to 0.6 per hour, $p_2$ to 0.4 per hour, $p_3$ to 150 per hour, $c_1$ to 80, $c_2$ to 35, $c_3$ to 25, $s_1$ to 24 per hour, $s_2$ to 70 per hour, $a_1$ to 1.5 per hour, $l_{cn}$ to 0.63 per hour, $l_{nc}$ to 0.15 per hour and $\theta_1$ and the corresponding rates for $S_2$ and $S_3$ to 2 per hour. The remaining seven parameters ($a_2$, $b_1$, $b_2$, $b_3$, $l_b$, $u_1$ and $u_2$) were fixed by requiring a steady state when no increased starvation signal was present, i.e. when $S_1$, $S_2$ or $S_3$ are 0.01.
