## Supplementary material for "Multi-signal regulation of the GSK-3β homolog Rim11 governs meiosis entry in yeast": Table S3

**Table S3. Oligo nucleotide sequences used**

| name | sequence | gene | notes |
|---|---|---|---|
| JK_IME2_qRT_F | CAGATTTTGGTTTGGCACGC | IME2 | |
| JK_IME2_qRT_R | TACAGTAACTTCCACCGCCA | IME2 | |
| FvWACT1Frt | gtaccaccatgttcccaggtatt | ACT1 | |
| FvWACT1Rrt | caagatagaaccaccaatccaga | ACT1 | |
| pUB415 | TGCCTCTTTAGGCGATTCGT | IME2 | Fig 6E, Fig S6 |
| pUB416 | GCTCGAACTTTTCCCGTGATT3 | IME2 | Fig 6E, Fig S6 |
| pUB2598 | GTACCACCATGTTCCCAGGTATT3 | ACT1 | Fig 6E, Fig S6 |
| pUB2599 | AGATGGACCACTTTCGTCGT3 | ACT1 | Fig 6E, Fig S6 |
