## Supplementary material for "Multi-signal regulation of the GSK-3β homolog Rim11 governs meiosis entry in yeast": Table S4

**Table S4. Plasmids used in this study**

| Plasmid No. | Name |
|---|---|
| FW_P770 | pNH605-RIM11-mNG |
| FW_P772 | pNH605-RIM11_S5AS8AS12A-mNG |
| FW_P779 | pNH605-RIM11Y199F-mNG |
| FW_P780 | pNH605-RIM11_NLS-mNG |
| FW_P730 | pNH604-pIME1-sfGFP-IME1-L321F |
| FW_P506 | pNH604-pIME1-sfGFP-IME1 |
